# Feedback control of Wnt signaling based on histidine cluster co-aggregation between Naked/NKD and Axin

**DOI:** 10.1101/2020.06.10.144725

**Authors:** Melissa Gammons, Miha Renko, Joshua E. Flack, Juliusz Mieszczanek, Mariann Bienz

## Abstract

Feedback control is a universal feature of cell signaling pathways. Naked/NKD is a widely conserved feedback regulator of Wnt signaling which controls animal development and tissue homeostasis. Naked/NKD destabilizes Dishevelled, which assembles Wnt signalosomes to inhibit the β-catenin destruction complex via recruitment of Axin. Here, we discover that the molecular mechanism underlying Naked/NKD function relies on its assembly into ultrastable decameric core aggregates via its conserved C-terminal histidine cluster (HisC). HisC aggregation is facilitated by Dishevelled and depends on accumulation of Naked/NKD during prolonged Wnt stimulation. Naked/NKD HisC cores co-aggregate with a conserved histidine cluster within Axin, to destabilize it along with Dishevelled, possibly via the autophagy receptor p62 which binds to HisC aggregates. Consistent with this, attenuated Wnt responses are observed in CRISPR-engineered flies and human epithelial cells whose HisC has been deleted. Thus, HisC aggregation by Naked/NKD provides context-dependent feedback control of prolonged Wnt responses.

## INTRODUCTION

Cells communicate via a handful of highly conserved signaling pathways to coordinate growth and differentiation. Signal responses are often mediated by signalosomes, dynamic protein condensates that are assembled transiently at the plasma membrane by hub proteins, to transduce incoming signals to the cytoplasm and nucleus of cells (Bienz, 2014; Case et al., 2019; Schaefer and Peifer, 2019). A well-studied hub protein is Dishevelled, a pivotal component within the ancient Wnt signaling pathway which controls cell specification during development and adult tissue homeostasis throughout the animal kingdom (Cadigan and Nusse, 1997; Clevers et al., 2014). Dishevelled transduces Wnt signals through canonical or non-canonical effector branches to elicit distinct cellular readouts, whereby the best-studied one is the β-catenin-dependent canonical branch which requires interaction of Dishevelled with Axin (Angers and Moon, 2009; Gammons and Bienz, 2018).

In order to transduce Wnt signals to β-catenin, Dishevelled assembles signalosomes by dynamic head-to-tail polymerization of its DIX domain (Schwarz-Romond et al., 2007). This enables it to bind to the DIX domain of Axin, which leads to inhibition of the associated kinases in the β-catenin destruction complex (Stamos et al., 2014) whose assembly, in the absence of signaling, depends on polymerization by the Axin DIX domain (Fiedler et al., 2011). In other words, the flow through the canonical Wnt signaling pathway is determined by the opposing activities of polymerizing Axin and Dishevelled and their mutual interaction via their DIX domains (Bienz, 2014): the polymerization of Axin promotes destabilization of β-catenin and therefore ensures quiescence of the pathway, while the polymerization of Dishevelled allows signalosome assembly and co-polymerization with Axin, which allows β-catenin to accumulate in the nucleus and engage in transcriptional activation of Wnt target genes (Gammons and Bienz, 2018).

Polymerization of Dishevelled and Axin is concentration-dependent (Schwarz-Romond et al., 2007), and occurs spontaneously if the cellular concentration of either of these proteins rises above a critical threshold. This undermines the physiological control of the signaling flux by Wnt proteins, with detrimental consequences for cells including lethality (e.g. Fiedler et al., 2011; Penton et al., 2002). It is therefore imperative that the levels of Dishevelled and Axin are tightly controlled in receiving cells. Indeed, Dishevelled and Axin are each recognized by specific ubiquitin E3 ligases that allow their individual targeting for proteasomal degradation (Angers et al., 2006; de Groot et al., 2014; Ji et al., 2017; Mund et al., 2015; Wei et al., 2012; Zhang et al., 2014; Zhang et al., 2011a). In addition, Dishevelled is also targeted for degradation by autophagy via association with the autophagy receptor p62 and the ubiquitin-like proteins LC3 or GABARAPL (Gao et al., 2010; Ma et al., 2015; Zhang et al., 2011b). Note that any mechanism earmarking whole signalosomes for degradation could result in either termination or maintenance of Wnt responses, depending on whether the Wnt signaling flux is limited in a given receiving cell by Wnt agonists such as Dishevelled, or by Wnt antagonists such as Axin.

One of the least-understood factors promoting the degradation of Dishevelled is the feedback regulator Naked/NKD. Naked Cuticle (Naked) was discovered in *Drosophila* as an essential Wnt-inducible antagonist of the canonical Wnt pathway (Zeng et al., 2000): loss of Naked causes hyperactivation of Wingless (fly Wnt) signaling in the embryo, causing late embryonic lethality, and although Naked is no longer required during larval development, its overexpression in larval tissues disrupts both canonical and non-canonical Wingless signaling (Rousset et al., 2001). Humans and mice encode two paralogs of Naked (Li et al., 2004; Wharton et al., 2001), and a third paralog was found in zebrafish (Schneider et al., 2010; Van Raay et al., 2007) (vertebrate Naked proteins are collectively referred to as NKD). As in fly embryos, expression of Nkd1 is induced by Wnt signaling in zebrafish embryos – in contrast to Nkd2 and Nkd3 which are ubiquitous and maternally provided (Schneider et al., 2010; Van Raay et al., 2007). Furthermore, overexpression of NKD paralogs can antagonize both canonical and non-canonical Wnt signal transduction in fish and frog embryos (Hu et al., 2010; Marsden et al., 2018; Van Raay et al., 2007; Yan et al., 2001), while depletion of Nkd1 and Nkd2 in early zebrafish embryos results in hyperactivation of canonical Wnt target genes. In addition, Nkd1 depletion produces non-canonical defects in left-right asymmetry signaling (Marsden et al., 2018; Schneider et al., 2010), exacerbates convergent extension defects under certain conditions (Van Raay et al., 2007) and sensitizes embryos to hyperactive Wnt signaling (Agonin and Van Raay, 2013). Curiously, in the light of these defects caused by Naked/NKD loss in fly and fish embryos, *Nkd* does not appear to be essential in mice: five double-knockout mice were bread from heterozygous parents, although these were born at sub-Mendelian ratios, and with subtle bone abnormalities reminiscent of *Axin2*-deficient mice (Zhang et al., 2007). Their fertility or longevity was not examined, and there was also a discrepancy with an earlier testis-related phenotype in *Nkd1* mutant mice (Li et al., 2005) that was not fully resolved. Therefore, while the available data are consistent with a partially redundant function of *Nkd* as a negative feedback regulator of canonical Wnt signaling in mice (Zhang et al., 2007), a systematic follow-up study is required for a definitive assessment of the physiological role of *Nkd* in this model organism. Notably, Naked/NKD is the only known intracellular feedback regulator of Wnt signaling that is conserved throughout the animal kingdom (**Fig. S1**), which argues strongly for its ubiquitous function and physiological relevance. Note also that feedback control is a wide-spread feature of signaling pathways and serves to canalize signal responses during development, and render them robust (Freeman, 2000).

Molecularly, Naked/NKD contains a single EF-hand, which binds to the PDZ domain of Dishevelled (Rousset et al., 2001; Rousset et al., 2002; Wharton et al., 2001; Yan et al., 2001). This results in destabilization of Dishevelled, apparently via the ubiquitin/proteasome system (Guo et al., 2009; Hu et al., 2010; Schneider et al., 2010). Indeed, Dishevelled was identified as a key target for downregulation by NKD (Rousset et al., 2001; Van Raay et al., 2007; Wharton et al., 2001; Yan et al., 2001), consistent with a role of Naked/NKD as an antagonist of canonical Wnt signaling. It could also explain why canonical and non-canonical signaling defects have been observed in flies and in vertebrate embryos upon overexpression of Naked/NKD (see above).

Here, we examine the molecular mechanism by which Nkd1 controls Wnt responses in human epithelial cells. This critically depends on a highly conserved histidine cluster (HisC) in its C-terminus which forms ultrastable decameric core aggregates *in vitro*. Nkd1 also forms HisC aggregates upon accumulation in Wnt-stimulated cells, promoted by its interaction with Dishevelled. Notably, these HisC aggregates bind selectively to Axin in vitro and in vivo, thereby promoting its destabilization in cells. We used CRISPR-engineering to generate human epithelial cell lines bearing specific HisC deletions of both NKD paralogs, which compromises their ability to sustain Wnt signal transduction to β-catenin and to destabilize Axin during prolonged Wnt stimulation. Similarly, CRISPR-engineered HisC deletion of *Drosophila nkd* results in embryonic defects reflecting reduced Wingless responses. These results indicate the physiological relevance of Naked/NKD HisC in flies and human cells, and reveal cellular contexts in which Naked/NKD acts as an agonist of Wnt signaling by promoting the destabilization of Axin via HisC aggregation. We also discover the autophagy p62 receptor as a HisC-dependent binding partner of Nkd1, implicating autophagy as the underlying mechanism.

## RESULTS

### The Nkd1 HisC is crucial for ternary complex formation with Axin and DVL2

We used co-overexpression assays in HEK293T cells, monitoring Wnt signaling with a co-transfected β-catenin-dependent transcriptional reporter (SuperTOP) (Veeman et al., 2003), to confirm that murine HA-Nkd1 reduces the signaling activity of co-overexpressed DVL2-GFP (a human Dishevelled paralog), but not of co-overexpressed β-catenin (**Fig. S2**). Previous work established that this downregulation depends on the EF-hand of Nkd1, the Dishevelled-binding domain (Rousset et al., 2001; Rousset et al., 2002; Wharton et al., 2001; Yan et al., 2001). In addition, HA-Nkd1 is also less active in downregulating β-catenin signaling if its C-terminal HisC is deleted (ΔHisC), despite the substantially higher expression levels of this deletion mutant (**Fig. 1A**). The latter is consistently observed, suggesting that HisC functions to destabilize Nkd1.

**Fig. 1.**
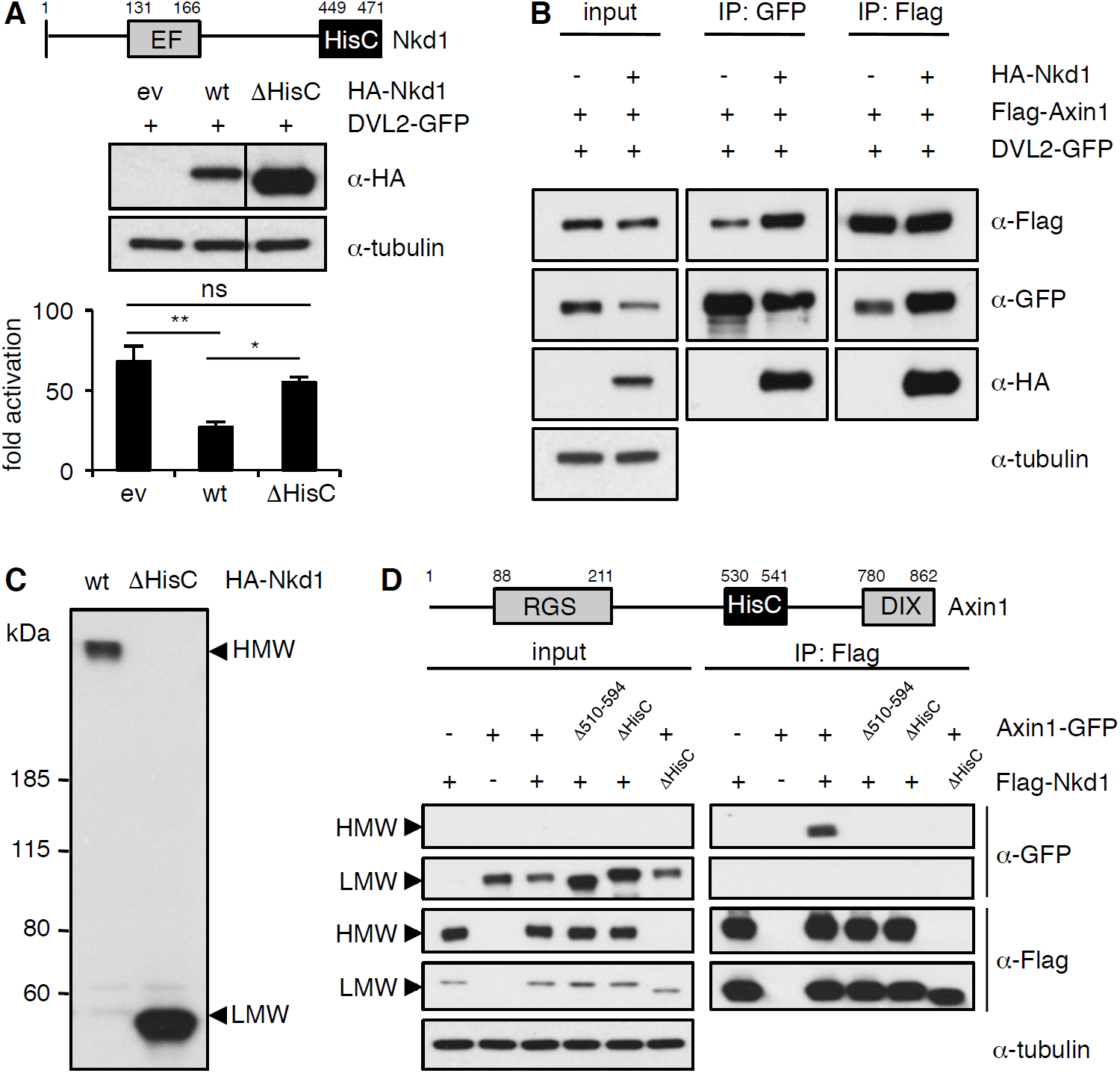
Nkd1 co-aggregates with Axin1 via highly conserved HisCs. (**A**) SuperTOP assays, monitoring the signaling activity of overexpressed DVL2-GFP upon co-expression with wt or ΔHisC HA-Nkd1 in transfected HEK293T cells (levels monitored by Western blot, *above*); error bars, SEM of >3 independent experiments; one-way ANOVA with multiple comparisons; * = p < 0.05, ** = p < 0.01. (**B**) CoIP assays between co-expressed proteins, as indicated above panels. (**C**) Western blot of wt and ΔHisC HA-Nkd1, revealing HisC-dependent HMW aggregates; *left*, positions of molecular weight markers (LMW, corresponding to monomeric Nkd1); in subsequent figures, only HMW and LMW regions are shown (see also supplementary information, for full-length gels). (**D**) CoIP assays between co-expressed proteins, as indicated above panels, monitoring ΔHisC-dependent co-aggregation of Flag-Nkd1 and Axin1-GFP (note that the amounts of transfected DNA has been adjusted in this experiment, to obtain comparable expression levels of different constructs). See also **Figs. S1-S3**.

Using co-immunoprecipitation (coIP) to monitor the association between co-expressed HA-Nkd1 and DVL2-GFP, we found that HA-Nkd1 also coIPs with Flag-Axin1 upon co-expression (**Fig. 1B**), as previously observed (Miller et al., 2009). This indicates that the three proteins can form a ternary complex. While conducting these coIP assays, we discovered that overexpressed Nkd1 forms high-molecular weight (HMW) aggregates that are resistant to boiling in sodium dodecyl sulphate (SDS), and whose formation critically depends on HisC (**Fig. 1C**). Furthermore, Axin1-GFP specifically co-aggregates with HMW but not LMW (low-molecular weight) Flag-Nkd1 via its own internal HisC (**Fig. 1D**). Thus, Nkd1 appears to be targeted to Axin-containing Wnt signalosomes via simultaneous binding to DVL and HisC-dependent co-aggregation with Axin.

### NKD1 HisC specifically co-aggregates with the internal HisC of Axin

Next, we tested whether recombinant HisC from NKD1 (identical sequence in mouse and human) would self-aggregate *in vitro*, by purifying lipoyl domain (Lip)-tagged protein (Lip-NKD-HisC) following expression in bacteria. Indeed, Lip-NKD-HisC forms stable aggregates that elute as a broad peak in the 250-450 kDa range (corresponding to 15-25-mers), as monitored by size exclusion chromatography (SEC) **(Fig. 2A)**. If the pH is lowered below ∼6.5, these aggregates begin to dissociate, and Lip-NKD-HisC is completely monomeric at pH 5 **(Fig. 2B)**. This sharp pH-dependence reflects the pKa value (∼6.0) of the imidazole side-chain of His: in acidic conditions (pH < 6), this side-chain is protonated and positively charged, and aggregation is thus blocked by electrostatic repulsion, whereas at higher pH values (pH > 6.2), the majority of His side-chains are uncharged, which is permissive for NKD-HisC core formation. Most probably, the majority of the His residues in Lip-NKD-HisC have to be deprotonated for aggregation (i.e. become uncharged, achieved at pH > pKa), while the protonation of a few critical His residues (pH < pKa) triggers disassembly of NKD-HisC aggregates owing to electrostatic repulsion between positively charged histidines.

**Fig. 2.**
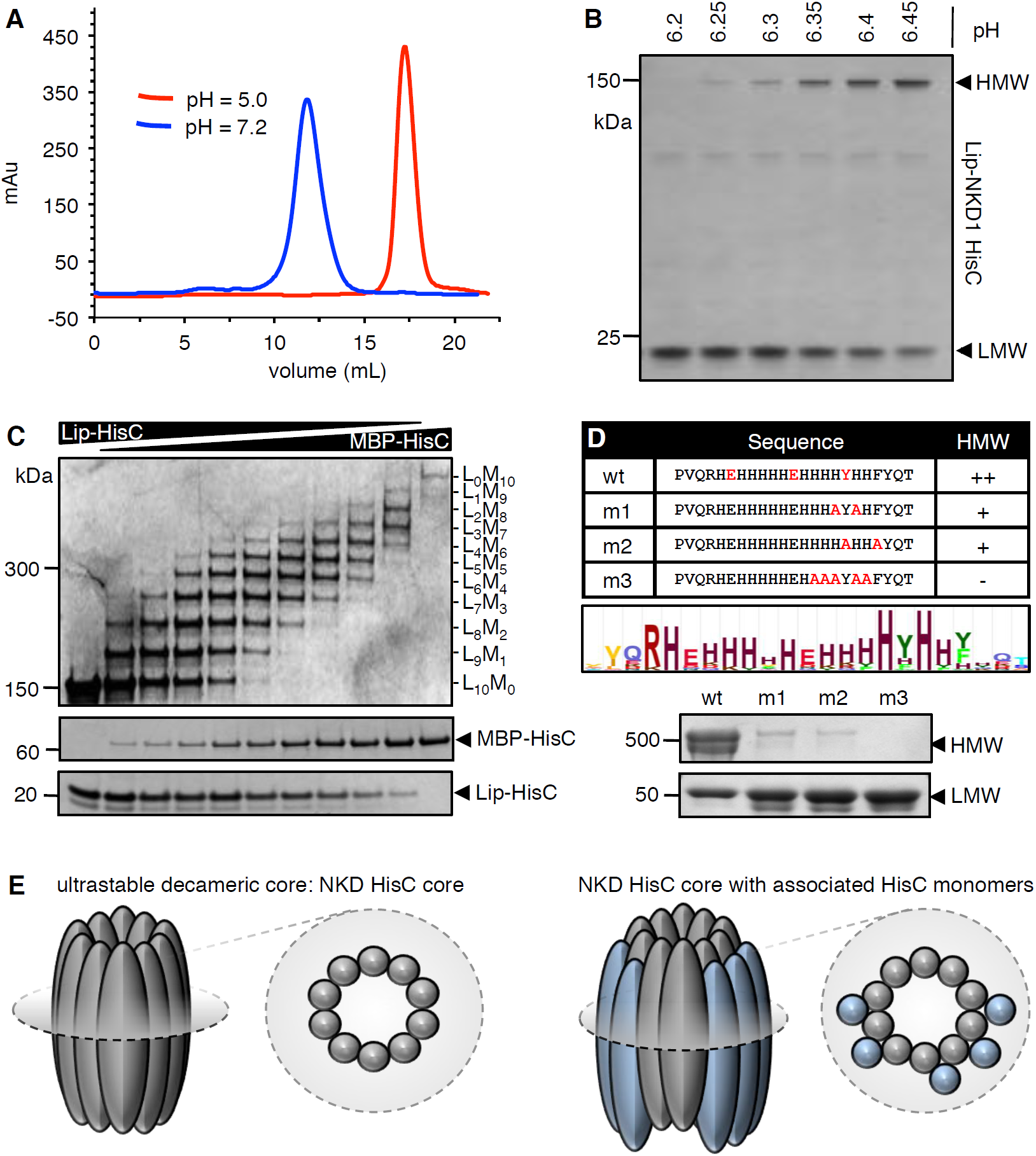
NKD1 HisC forms decameric core aggregates. (**A, B**) pH-dependent aggregation of purified recombinant Lip-NKD-HisC, monitored by (**A**) SEC or (**B**) PAGE. (**C**) Co-aggregation assays after mixing Lip-NKD-HisC and MBP-NKD-HisC at different ratios, revealing a decameric core (note that the self-aggregation of MBP-NKD-HisC is considerably less efficient than that of Lip-NKD-HisC, presumably owing to the large size of the MBP tag). (**D**) Mutational analysis, revealing that aggregation depends on conserved aromatic and charged residues within HisC. (**E**) Cartoon of NKD HisC core and associated HisC monomers.

If assayed by Polyacryl-Amide Gel Electrophoresis (PAGE) under denaturating and reducing conditions (i.e. after boiling in 1% w/v SDS and 5% v/v mercapto-ethanol), an ultra-stable NKD-HisC core of ∼150 kDa is observed. This core was sized more accurately by mixing monomeric Lip-NKD-HisC with monomeric NKD-HisC bearing a maltose-binding protein tag (MBP-NKD-HisC) at different ratios at acidic pH, and by increasing the pH subsequently, to allow aggregation. Because of the different molecular masses of these tags, we observe a discrete laddering on PAGE, revealing that the stable NKD-HisC core consists of a 10-mer (**Fig. 2C**). Notably, the formation of NKD-HisC aggregates not only depends on the most conserved histidines, but also on two aromatic residues within HisC that are highly conserved amongst Naked/NKD orthologs **(Fig. S1)**: double- or quintuple point-substitutions of the most conserved histidine or aromatic residues (m1-m3) strongly reduce aggregation of recombinant Lip-NKD-HisC (**Fig. 2D)**. Our results indicate that NKD-HisC aggregates into a decameric ultra-stable core (called HisC core below) that can be decorated by additional NKD-HisC monomers that are less firmly associated (**Fig. 2E**).

We also tested full-length HA-Nkd1 bearing these point-substitutions in transfected HEK293T cells, for their ability to form HisC aggregates and to block β-catenin signaling by co-expressed Flag-DVL2. Indeed, the two double-point mutations (m1, m2) attenuate aggregation of HA-Nkd1 in cells, particularly noticeable when co-expressed with Flag-DVL2, while the quintuple mutation (m3) is almost as strong as ΔHisC in blocking aggregation (**Fig. S3**). Notably, each of the three mutations also stabilizes HA-Nkd1, with m3 being almost as potent as ΔHisC. However, despite their significantly increased levels, none of the three mutants is as active as wt HA-Nkd1 in antagonizing Flag-DVL2 signaling: m1 reduces signaling only partially, while m2 and m3 abolish the activity of HA-Nkd1 in antagonizing Flag-DVL2 signaling (**Fig. S3**). Therefore, these substitutions of highly conserved residues within HisC behave as expected from our in vitro results with recombinant HisC protein. They provide strong evidence that the HisC-dependent aggregation observed in vitro is required for the activity of Nkd1 in antagonizing DVL2 signaling in cells.

His-rich sequences are relatively rare in the genome. We searched the human genome for proteins exhibiting a minimum of 8 His residues within a 13 amino acid stretch, which identified 74 proteins (containing 78 HisC sequences; **Table S1**). Most of these are nuclear proteins and thus unlikely targets for Naked/NKD, which is anchored at the plasma membrane by its N-terminus via myristoylation (Chan et al., 2007; Hu et al., 2010). The non-nuclear set contains Axin1 and Prickle3 (a non-canonical Wnt signaling component) as well as a small number of additional proteins (**Fig. 3A**). We tested a subset of these HisC (tagged with MBP) for *in vitro* co-aggregation, and found that MBP-Axin1-HisC exhibits the most pronounced co-aggregation with Lip-NKD-HisC (<5 molecules per core, at equivalent ratio; **Fig. 3B)**. Axin2 also exhibits an internal HisC, which co-aggregates with Lip-NKD-HisC (<4 molecules per core, at equivalent ratio), although this seems insufficient for its co-aggregation with ultrastable Nkd1 HisC cores in cells (**Fig. S4**). The MBP-HisCs from the remaining tested HisC-containing proteins co-aggregate poorly, or not at all, with Lip-NKD-HisC (**Fig. S5**). Given the sequence conservation of the HisC in Axin orthologs, the specificity and avidity of co-aggregation between Axin1-HisC and NKD1 HisC cores in vitro and the co-aggregation between Axin1 and Nkd1 in cells, it seems likely that Axin1 is the primary if not only physiological target for NKD co-aggregation within the human genome.

**Fig. 3.**
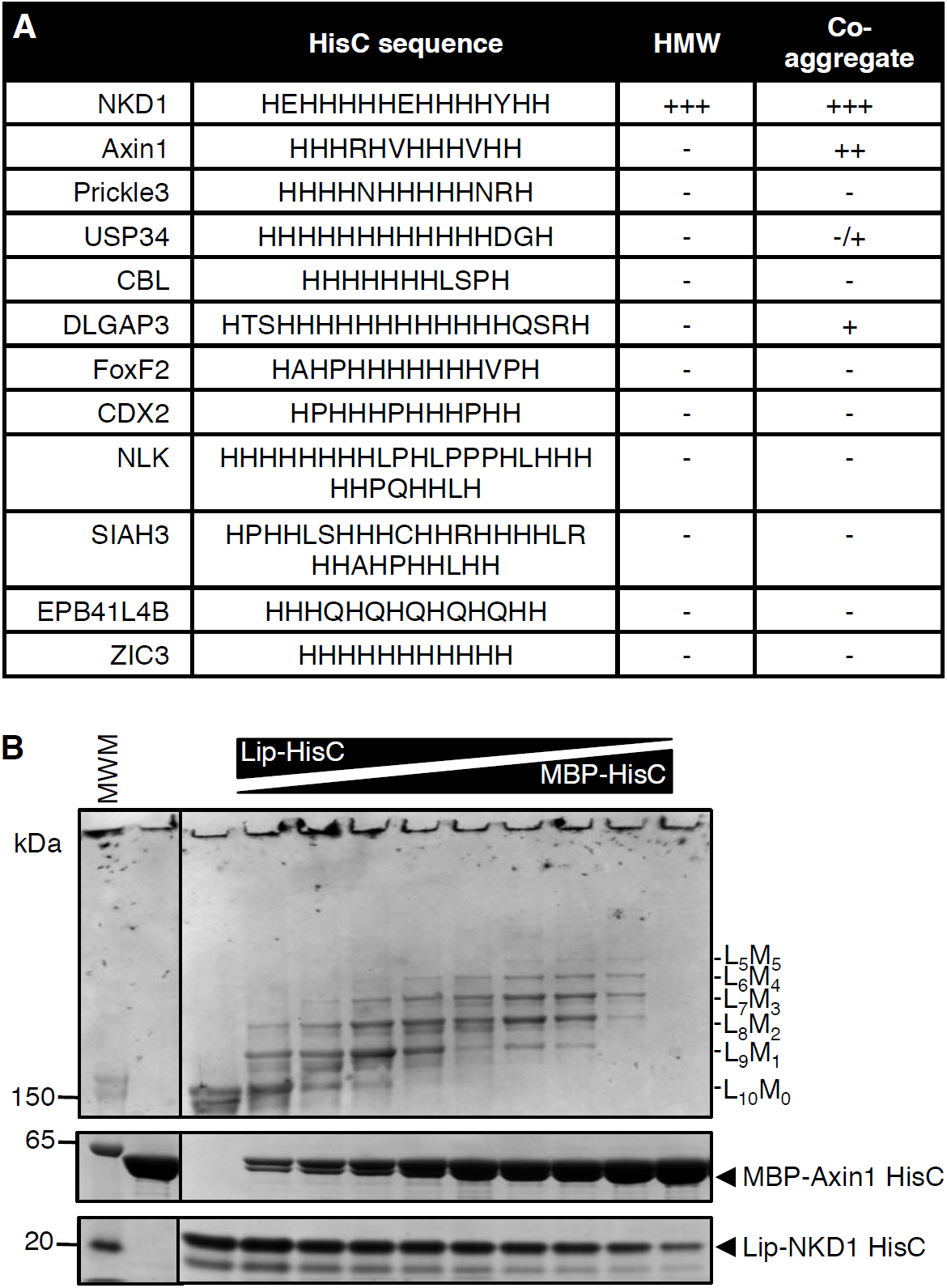
NKD1 HisC co-aggregates with Axin1 HisC. (**A**) Selected HisC-containing proteins in the human genome (see also text) and their ability to co-aggregate with NKD1 HisC. (**B**) Co-aggregation assays after mixing Lip-NKD-HisC and MBP-Axin1-HisC at different ratios (MWM, molecular weight markers). See also **Figs. S4 & S5, Table S1**.

### Dishevelled promotes co-aggregation between Nkd1 and Axin1

The level of SDS-resistant HMW Nkd1 aggregates clearly depends on the level of Nkd1 overexpression, indicating that HisC aggregation is concentration-dependent (**Fig. 4A**; note that this concentration dependence of HisC aggregation explains why the ratios between HMW and LMW Nkd1 vary somewhat between experiments, depending on the transfection efficiency). We therefore asked whether Wnt stimulation (which increases Nkd1 expression; Van Raay et al., 2007; Yan et al., 2001) would promote Nkd1 aggregation, which is indeed the case (**Fig. 4B**). In addition, Dishevelled stimulates efficient Nkd1 aggregation since the formation of HMW aggregates is much reduced in DVL null cells (lacking all three DVL paralog; Gammons et al., 2016b) (**Fig. 4C**). Testing a panel of DVL2-GFP mutants (Gammons et al., 2016a) for their ability to restore formation of HMW aggregates upon re-expression in these DVL null cells, we found that polymerization-deficient mutants or DVL2 lacking its DIX domain were most active, while DVL2 mutants whose DEP-dependent dimerization is blocked (E499G or G436P) were least active (**Fig. 4D; Fig. S6**). This indicates that the formation of stable DVL2 dimers formed by DEP domain swapping is crucial for Nkd1 aggregation. As expected, restoration of HMW aggregates also depends on the PDZ domain, the NKD-binding domain of DVL2 (Rousset et al., 2001; Wharton et al., 2001; Yan et al., 2001), but is hardly affected by K446M which blocks binding of the DEP domain to the Frizzled receptor (**Fig. 4D**).

**Fig. 4.**
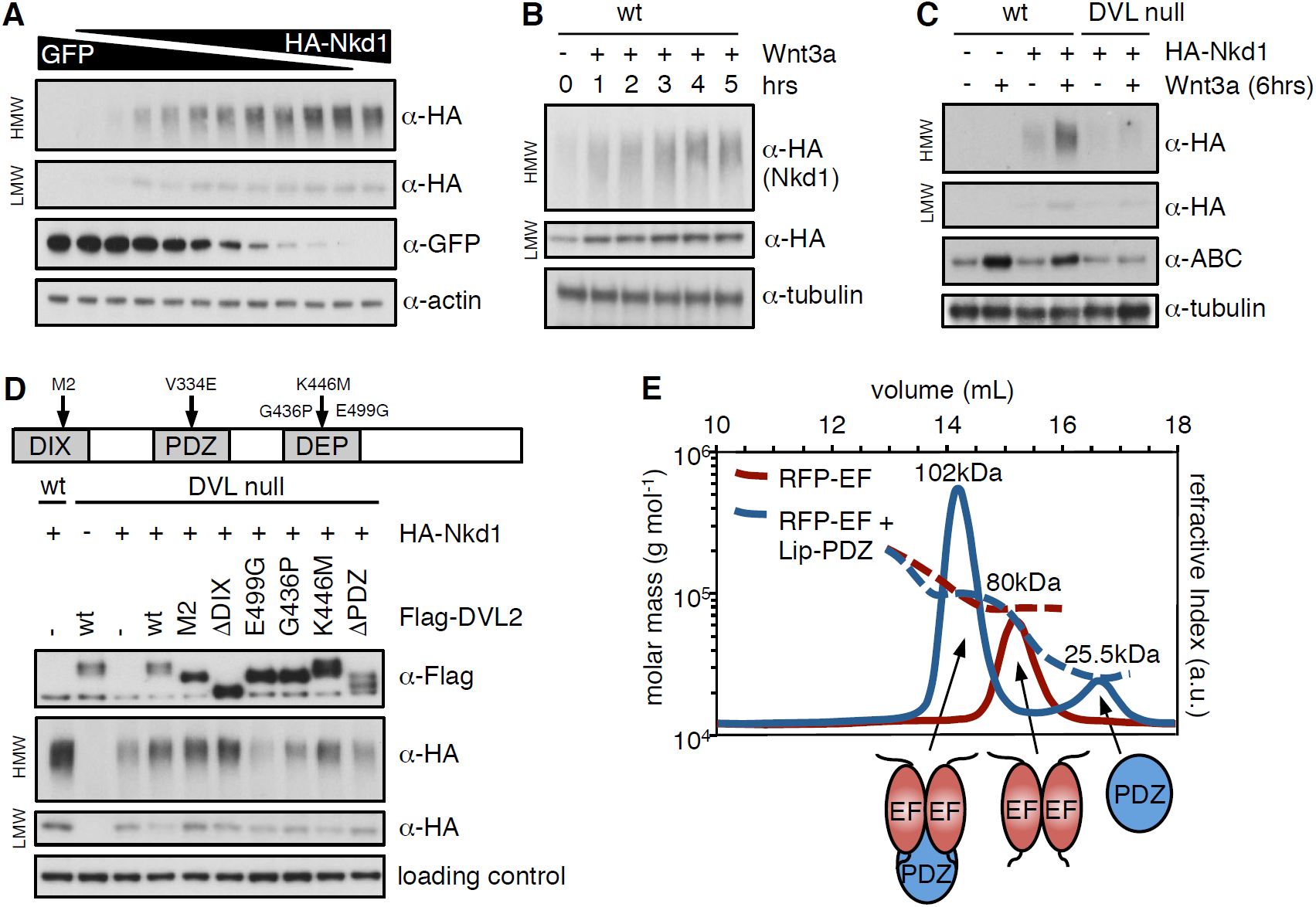
HisC aggregation is promoted by DVL2. (**A-D**) Western blots monitoring the formation of HisC aggregates dependent on (**A**) Nkd1 concentration, (**B**) duration of Wnt stimulation, (**C**) DVL or (**D**) complementation of DVL null cells by re-expression of wt or mutant DVL2 (E499G, G436P block DVL2 dimerization; K446M blocks Frizzled receptor binding) (Gammons et al., 2016a). (**E**) SEC-MALS, monitoring complex formation between purified recombinant RFP-NKD1-EF and Lip-DVL2-PDZ (as indicated by cartoons); numbers correspond to Mr values determined by MALS (expected: RFP-NKD1-EF dimer, 71.8 kDa; Lip-DVL2-PDZ, 22.0 kDa; complex, 93.8 kDa). See also **Fig. S6**.

To confirm that the DVL2 PDZ domain binds directly to the Nkd1 EF-hand, we used SEC followed by Multi-Angle Light Scattering (SEC-MALS) of purified recombinant domains to show that a stable complex forms between them (**Fig. 4E**). Using isothermal titration calorimetry (ITC), we estimated a binding affinity in the low-micromolar range (*K*_d_ 3.2 ± 0.2μM), which depends on the integrity of the PDZ cleft as demonstrated by the failure of the EF-hand to bind to the V334E cleft mutant (**Fig. S6**) whose binding to cognate substrate motifs is abolished (Gammons et al., 2016b). As expected, this cleft mutant also fails to restore Nkd1 HMW aggregates in DVL null cells (**Fig. S6**). Interestingly, the molecular mass of the EF-hand-PDZ complex indicates that the PDZ domain binds to an EF-hand dimer (**Fig. 4E**). Taken together with our previous results (**Fig. 4D**), this implies that a single DVL2 dimer can bind to four molecules of Nkd1, resulting in a considerable increase of its local concentration, which may provide the trigger for the formation of HisC core aggregates.

### NKD HisC is required for Wnt-induced Axin destabilization

Our in vitro and cell-based assays uncovered the striking activity of the NKD HisC to form hyperstable aggregates and co-aggregates with Axin (**Figs. 1-3, S3-S5**). However, these assays are based on recombinant NKD HisC or on overexpression of Nkd1 in cells. We therefore decided to use CRISPR to engineer specific HisC deletions in both NKD paralogs in HEK293T cells (NKDΔHisC), to test the physiological relevance of these histidine clusters and their interaction with Axin. As a comparison, we also generated double-knockout cells (NKD null cells), to assess the function of NKD on Wnt signaling and Axin stability (**Fig. S7**).

Somewhat to our surprise, NKD null cells show a normal response to Wnt3a during the first ∼3 hours (hrs) (**Fig. 5A, B**), with no detectable hyperactivation of β-catenin-dependent transcription. However, by ∼6 hrs of Wnt3a stimulation, their signaling response is noticeably attenuated and begins to plateau, unlike the response of parental control cells which continues to rise, reaching at least three times the levels compared to that of the NKD null cells by 12 hrs (**Fig. 5A**). This is observed in two independently isolated NKD null lines, based on different targeting events (**Fig. S7**). Similarly, the Wnt response is also significantly reduced from 6 hrs onwards in two independently isolated NKDΔHisC lines, although this reduction is less pronounced than in NKD null cells (**Fig. 5A**). Similar trends are observable if the levels of endogenous activated β-catenin are monitored with an antibody specific for the activated form (α-ABC; **Figs. 5B & S8**). These results provide clear evidence that NKD functions as a positive regulator of Wnt signaling in these human epithelial cells, and reveal the functional importance of its histidine cluster in the maintenance of Wnt responses during prolonged Wnt stimulation.

**Fig. 5.**
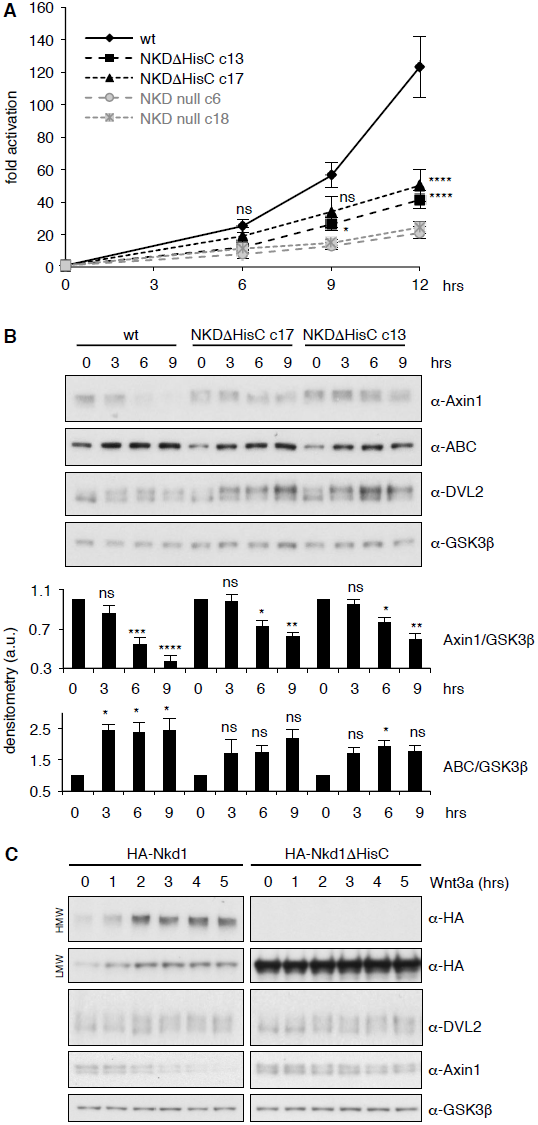
NKD is required for sustained Wnt signaling. (**A**) SuperTOP assays, monitoring the signaling response of NKDΔHisC, NKD null mutant or control cells after 6 hrs of Wnt3a stimulation (two independently isolated lines each, see also **Fig. S7**); error bars, SEM of >3 independent experiments; two-way ANOVA with multiple comparisons (for c13 and c17; see **Fig. S8** for null lines); * = p < 0.05, **** = p < 0.0001. (**B**) Representative Western blots monitoring expression levels of endogenous proteins in two independent NKDΔHisC lines following 0-9 hrs of Wnt stimulation (as indicated), quantified by densitometry of >3 independent experiments relative to GSK3β as internal control (a.u., arbitrary units); error bars, SEM of >3 independent experiments; two-way ANOVA with multiple comparisons; ^*^ = p < 0.05, ^**^ = p < 0.01, ^***^ = p < 0.001, ^****^ = p < 0.0001, ns = not significant. For the corresponding analysis of NKD null cell lines, see **Fig. S8**. (**C**) Western blots monitoring the effects of overexpressed wt or ΔHisC HA-Nkd1 on the levels of endogenous proteins (indicated on the right; GSK3β, internal control) after 0-5 hrs of Wnt stimulation. See also **Figs. S7 & S8**.

The reduction of β-catenin signaling in NKD-deficient cells coincides roughly with the half-time (t1/2) of the known Wnt-induced destabilization of Axin1 (Ji et al., 2017; Yamamoto et al., 1999). Indeed, while the levels of endogenous Axin1 gradually decrease to approximately half within the first 6 hrs after Wnt stimulation in parental HEK293T cells, they decrease less in NKDΔHisC cells (**Fig. 5B**), and there is no statistically significant reduction of the Axin1 levels in NKD null cells (**Fig. S8**). Similarly, the levels of endogenous DVL2 are elevated in NKDΔHisC (**Fig. 5B**) and NKD null cells (**Fig. S8**), most noticeable for the Wnt-induced phosphorylated form. Thus, Axin1 and DVL2 are both stabilized during prolonged Wnt stimulation in cells without functional NKD, indicating that NKD functions to target both signalosome components for degradation in these cells. Its requirement for maintaining β-catenin responses can readily be explained by its destabilizing effect on Axin1 but not on DVL2 (see also Discussion), which argues that Axin1 rather than DVL2 is rate-limiting for a sustained Wnt response of these cells.

To test this further, we asked whether Nkd1 might accelerate the Wnt-induced degradation of Axin1 and DVL2 upon overexpression. Indeed, overexpressed Nkd1 but not Nkd1ΔHisC (incapable of forming HMW aggregates) accelerates the destabilization of endogenous Axin1 upon Wnt stimulation, but does not affect the levels of endogenous DVL2 (**Fig. 5C**). This provides further support for the critical role of the Nkd1 HisC in the Wnt-induced destabilization of Axin1, and suggests that Axin1 rather than DVL2 is the relevant target of NKD in persistently Wnt-stimulated HEK293T cells.

### The phenotype of *Drosophila nkd*^*Δhis*^ mutants implies attenuated Wingless signaling

Given the unexpected reduction of β-catenin signaling in NKD-deficient human cells, we decided to revisit the *nkd* mutant phenotype in *Drosophila* embryos whose signature ‘naked cuticle’ in a putative null allele (*nkd*^*7H16*^) indicates hyperactive Wingless signaling (Zeng et al., 2000) (**Fig. 6A, B**). However, these authors noted a caveat regarding a possible genetic interaction with *h*^*1*^, a recessive marker born on the original chromosome (Nusslein-Volhard et al., 1984). Indeed, the cuticle phenotype of another strong allele (*nkd*^*7E89*^) is far milder upon removal of the *h*^*1*^ allele (Waldrop et al., 2006) (**Fig. 6C, D**).

**Fig. 6.**
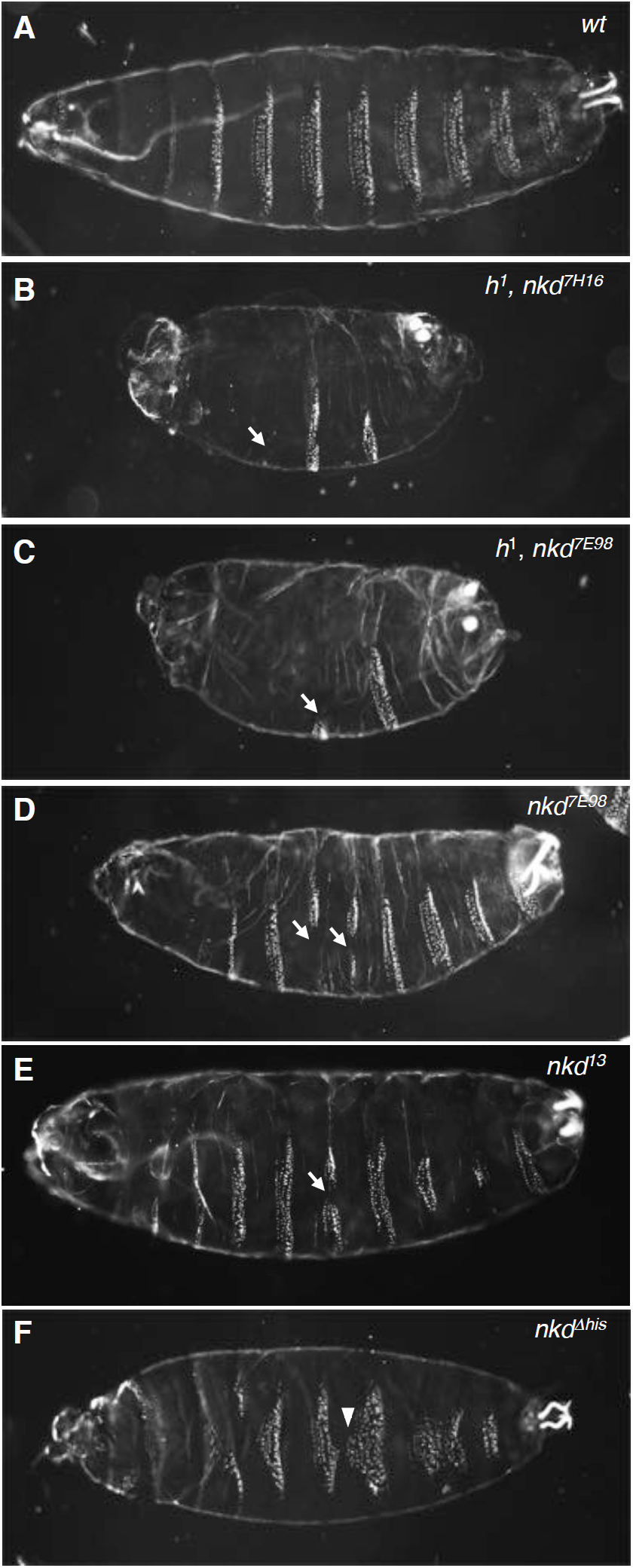
Allele-dependent Wg signaling defects in *nkd* mutant *Drosophila* embryos. Dark-field images of ventral cuticles of (**A**) wt and (**B-F**) homozygous mutant embryos, as indicated in panels (*nkd*^*7H16*^, *nkd*^*7E98*^, pre-existing alleles; *nkd*^*13*^, *nkd*^*Δhis*^ CRISPR-engineered alleles), showing regions of excess naked cuticle (arrows) signifying ectopic Wg signaling. Note that the ‘near-naked’ phenotype is only seen in embryos bearing *h*^*1*^ (**B, C**); excess naked cuticle regions are seen infrequently in null mutant embryos without *h*^*1*^ (**D, E**), or not at all in *nkd*^*Δhis*^ embryos bearing a HisC deletion (**F**) whose most penetrant phenotype are excess small denticles (arrowheads) signifying reduced Wg signaling. See also **Fig. S9**.

To determine the true *nkd* null phenotype, we generated a CRISPR-engineered null allele (*nkd*^*13*^) truncating Naked at codon 172 upstream of its EF-hand (**Fig. S9**). Freshly hatched *Drosophila* larvae show 8 ventral abdominal denticle belts comprising 6 denticle rows each (**Fig. 6A**), whereby the denticles in each row exhibit a characteristic shape and orientation, specified by row-specific selector gene products (e.g. Engrailed, En) and by positional signals such as Wingless (Wg), Hedgehog and Spitz (an Epidermal Growth Factor-like factor activated by the transmembrane protease Rhomboid) (Hatini and DiNardo, 2001; Martinez Arias et al., 1988; O’Keefe et al., 1997; Szuts et al., 1997) (**Fig. 7A**). As expected, homozygous *nkd*^*13*^ embryos exhibit stretches of excess naked cuticle, however, these mutant embryos also show near-normal denticle belts that are uninterrupted by naked patches (**Fig. 6E**), more so than the mutant embryos bearing *h*^*1*^ (**Fig. 6B, C**). Notably, patches of missing denticles in *nkd*^*13*^ embryos are predominantly observed in rows 3-7 (**Fig. 7B**), and seem to correlate with the sporadic patches of ectopic Wg expression (Martinez Arias et al., 1988) (**Fig. S10**), owing to a failure of En and its *rhomboid* target gene to repress *wg* posteriorly to their own expression domains (Zeng et al., 2000; O’Keefe et al., 1997; Szuts et al., 1997) (**Fig. 7A**).

**Fig. 7.**
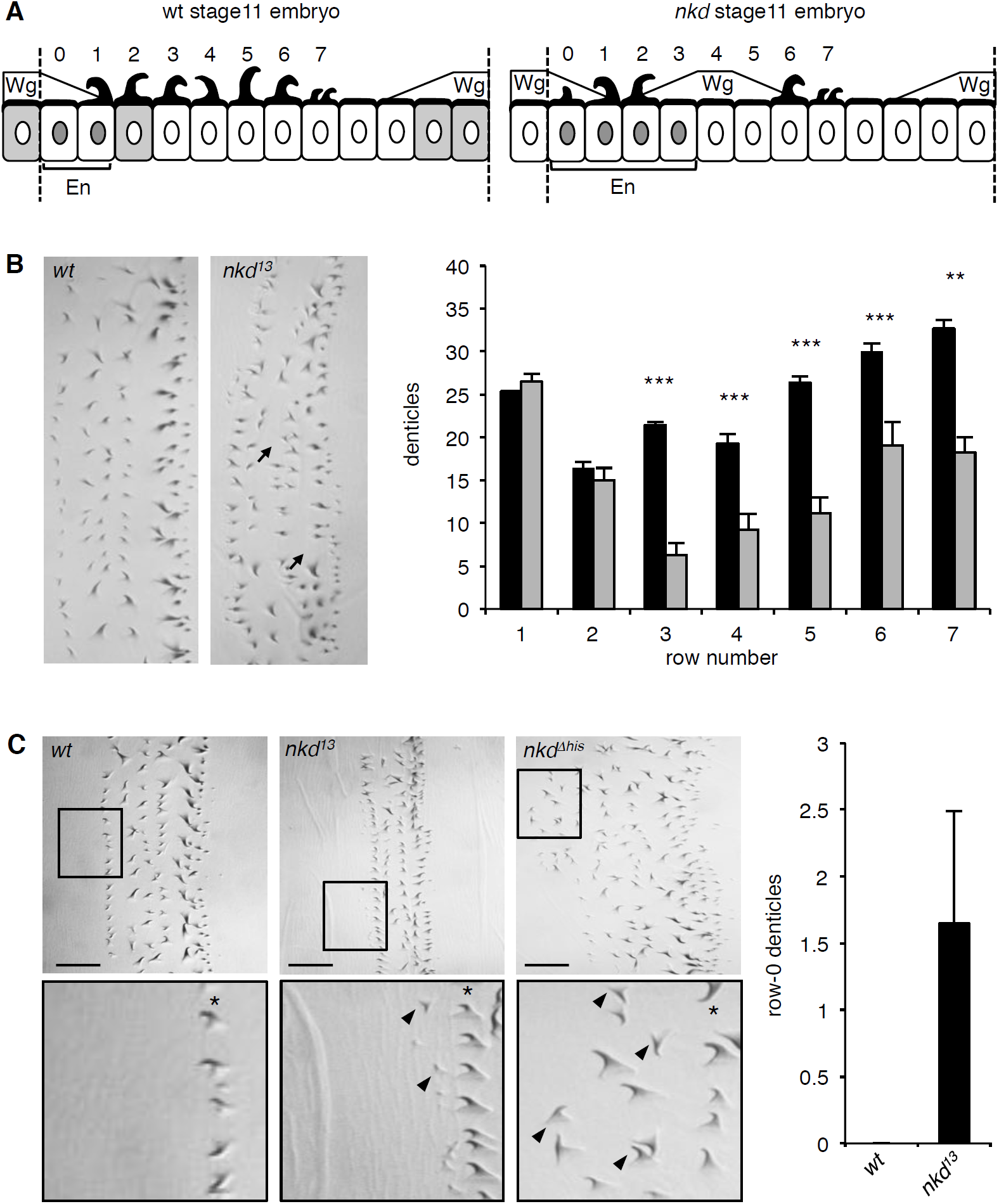
Quantification of context-dependent Wg signaling defects in embryos bearing *CRISPR alleles*. (**A**) Cartoons of row-specific ventral denticles relative to expression domains of Wg, En and Nkd in stage 11 wt (left) and *nkd*^*13*^ null mutant (right) embryos; numbers indicate denticle rows; see also text. (**B**) *Left*, examples of abdominal denticle belts in wt and *nkd*^*13*^ homozygous embryos, with arrows pointing to missing denticles; *right*, quantitation of numbers of denticles per row in wt (black bars) or *nkd*^*13*^ null mutant (white bars) embryos (n=10 near-normal denticle belts); Student’s unpaired t-test; ** = p < 0.01, *** = p < 0.001. (**C**) *Left, nkd*^*13*^ and *nkd*^*Δhis*^ homozygous but not wt embryos exhibit excess row-0 denticles (arrowheads; asterisks mark row-1); *right*, quantitation of excess row-0 denticles in *nkd*^*13*^ mutants (n=20 near-normal denticle belts). See also **Figs. S10-S12**.

On close examination of the near-normal denticle belts in *nkd*^*13*^ embryos, we noticed ectopic small denticles anteriorly to these belts (**Fig. 7C**), i.e. in row 0 (**Fig. 7A**). This phenotype, observed in ∼20% of near-normal denticle belts of *nkd*^*13*^ homozygotes (**Fig. 7C**), signifies a reduced Wg response in this part of the segment. This is not a CRISPR off-target effect since we also observe excess row-0 denticles in various transheterozygous embryos, including transheterozygotes of *nkd*^*13*^ and *h*^*1*^ *nkd*^*7H16*^ and, occasionally, in the rare near-normal denticle belts of *h*^*1*^ *nkd*^*7H16*^ homozygotes (**Fig. S11**). These excess row-0 denticles are consistent with the phenotype in NKD null-mutant human cells (**Fig. 5A**), and likely to reflect a failure to maintain Wg responses through late embryonic stages (a condition that causes excess small denticles since reduced Wg signaling is permissive for denticle specification by Rhomboid and its Spitz substrate, which stimulates EGF receptor signaling; O’Keefe et al., 1997; Szuts et al., 1997). We also examined the expression of a Wg-responsive reporter from the *Ultrabithorax* gene (*UbxB.lacZ*) (Thuringer et al., 1993) as a direct readout of Wg signaling in the middle midgut of late-stage embryos (Riese et al., 1997), and found this to be reduced consistently in *nkd*^*13*^ mutant embryos (**Fig. S12**). This further underscores the notion that Naked functions in late embryonic stages to maintain Wg signaling responses.

Given our results from human cells implicating the NKD HisC in maintaining prolonged Wnt responses (**Fig. 5A**), we asked whether this might also be true for the HisC of *Drosophila* Naked. We therefore generated a deletion of the C-terminal HisC (*nkd*^*Δhis*^) by CRISPR engineering (**Fig. S9**). Homozygous *nkd* ^*Δhis*^ flies can be obtained, however ∼20% of their offspring die as late-stage embryos. Each of these embryos shows several abnormal denticle belts with disordered denticle rows, misoriented denticles as well as excess small denticles in their anterior regions (**Figs. 6F & 7C**), most likely including excess row-0 denticles since these can also be observed in transheterozygous *nkd* ^*Δhis*^/*nkd*^*13*^ embryos (**Fig. S11**). It thus appears that these excess anterior denticles in *nkd* ^*Δhis*^ mutant embryos parallels the Wnt signaling defect of the NKDΔHis human cells (**Fig. 5A**), implicating the C-terminal HisC of *Drosophila* Naked in the maintenance of the Wg response throughout late embryonic stages.

### Nkd1 HisC aggregates associate with the p62 autophagy receptor

To identify potential effectors of Naked/NKD in maintaining prolonged Wnt responses, we used a BioID proximity-labeling approach, tagging Nkd1 with the promiscuous biotin ligase BirA* (Roux et al., 2012) internally (within its non-conserved low-complexity linker sequences), to avoid interfering with its N-terminal myristoylation (Hu et al., 2010) or with the function of its C-terminal HisC. We confirmed that Nkd1-BirA* is as active as HA-Nkd1 in suppressing DVL2 signaling (J. E. F., thesis). We then introduced this bait as an inducible transgene into a specific genomic location of Flp-In-T-REx-293 cells to keep its expression low, as previously described (van Tienen et al., 2017). As controls, we used mutant Nkd1-BirA* versions bearing a point mutation in their N-terminal myristoylation site (G2A), or a deletion of their HisC (ΔHisC; **Fig. 8A**). Amongst the hits from this approach, we expected to identify DVL and Axin paralogs as direct Nkd1-binding proteins, but also bystanders or ‘vicinal’ proteins (Roux et al., 2012) that bind to, or are closely associated with, DVL or Axin. We considered hits as specific only if the spectral counts obtained with wt Nkd1-BirA* were >5-fold higher than those obtained with GFP-BirA*.

**Fig. 8.**
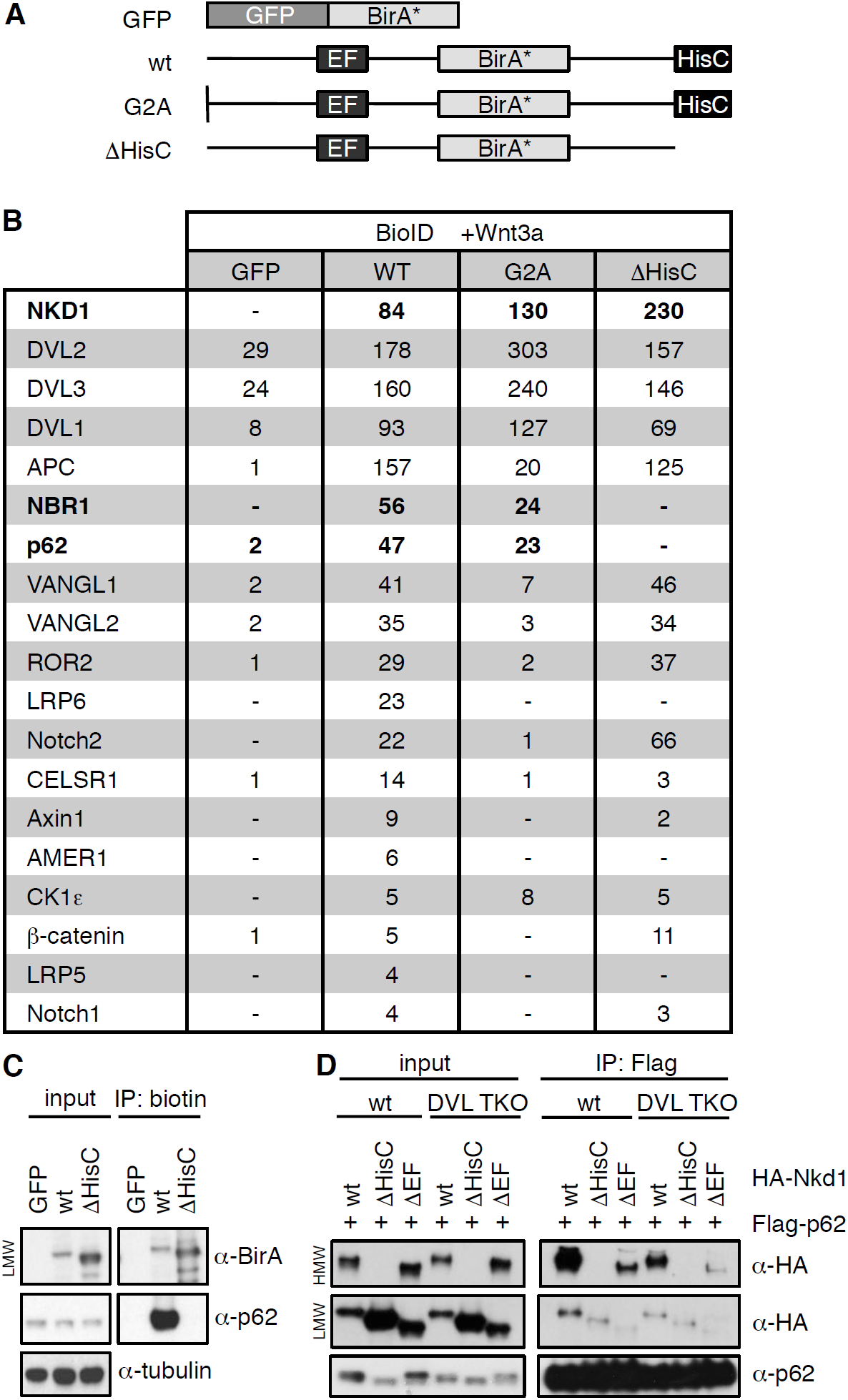
p62 is a HisC-dependent interactor of Nkd1. (**A**) Cartoons of BirA* baits used for BioID proximity labeling. (**B**) Selected BioID hits obtained with wt or mutant Nkd1-BirA* (numbers correspond to unweighted spectral counts >95% probability; for full list, see **Table S2**); note that APC is large (∼311 kDa) and relatively abundant, explaining the high numbers of spectral counts. (**C**) BioID proximity labeling assays, monitoring association between endogenous p62 and Nkd1-BirA* or Nkd1ΔHisC-BirA* (only LMW Nkd1-BirA* is shown since HMW Nkd1-BirA* is too large to be detectable on blots). (**D**) CoIP assays between co-expressed proteins, as indicated above panels, monitoring association between HA-p62 and wt or mutant HA-Nkd1. See also **Table S2.**

Cells expressing BirA* baits were stimulated with Wnt3a for 12 hrs while also being incubated with biotin, to label bait-interacting proteins. As expected, we found DVL1-3 amongst our top hits (**Fig. 8B & Table S2**). We also found components of the β-catenin destruction complex, namely Axin1 itself and its key binding partner, the APC tumour suppressor, as well as AMER1 and β-catenin, suggesting that the entire complex is associated with Nkd1 HisC aggregates in Wnt-stimulated cells. Also on our list were the Wnt co-receptors LRP5 and LRP6 whose cytoplasmic tails form stable complexes with Axin upon Dishevelled-dependent phosphorylation (Bilic et al., 2007; Tamai et al., 2004). Our myristoylation-dependent hits include the non-canonical Wnt signaling components VANGL1, VANGL2, CELSR1 and ROR2, as well as Notch1 and Notch2 (physiological substrates of Dishevelled-activated NEDD4 family ligases) (Mund et al., 2015) (**Fig. 8B & Table S2**). Strikingly, our top HisC-dependent hits are the two main autophagy receptors p62 and NBR1 (Kirkin et al., 2009). We used BioID proximity labeling (BioIP) to confirm that Nkd1-BirA* binds to endogenous p62 in a HisC-dependent fashion (**Fig. 8C**).

Dishevelled is a known substrate for selective autophagy, and is targeted to autophagosomes via association with p62, LC3 and GABARAPL and, ultimately, to lysosomes for degradation (Gao et al., 2010; Ma et al., 2015; Zhang et al., 2011b). However, the association between Nkd1 and p62 is independent of Dishevelled since the two proteins coIP as efficiently in DVL null cells as in control HEK293T cells (**Fig. 8D**). We used internal deletions to map the p62-interacting domain of Nkd1 to its EF-hand (**Fig. 8D**), which indicates that this EF-hand can bind Dishevelled and/or p62. As expected from our BioID screen, co-expressed Flag-p62 shows a strong preference for binding to HMW HA-Nkd1 (**Fig. 8D**). These results suggest that p62 may bind to Nkd1 HisC aggregates and Nkd1-Axin1 co-aggregates, to target these for autophagy-dependent degradation (see Discussion).

## DISCUSSION

From numerous studies in *Drosophila* and vertebrates, a picture has emerged of Naked/NKD as a widespread Wnt-inducible signaling component that downregulates canonical and non-canonical Wnt responses by binding to and destabilizing Dishevelled (see Introduction). Our results from CRISPR-engineered null mutants in *Drosophila* and human epithelial cells lead to a revised picture which also includes positive regulatory effects of Naked/NKD on canonical Wnt signaling, likely mediated by its binding to and destabilizing Axin during prolonged Wnt stimulation. Therefore, whether Naked/NKD antagonizes or promotes Wnt signaling appears to depend on whether Dishevelled or Axin is limiting for Wnt responses in any given cell. This context-dependence of Naked/NKD renders it a versatile feed-back regulator of the pathway, and may also explain why it is dispensable in some cells. Feedback control has long been recognised as a wide-spread mechanistic principle in cellular signaling that leads to canalization and robustness of development (Freeman, 2000).

Both negative and positive regulatory effects of Naked/NKD on Wnt signaling are mediated by its activity to destabilize its interaction partners. We discovered a rather unusual molecular mechanism underlying this destabilization, which depends on ultrastable aggregation of Naked/NKD via a highly conserved cluster of histidines in the Naked/NKD C-terminus. NKD HisC core aggregates are decameric, and their finite size implies a well-defined structure. Consistent with this, they co-aggregate specifically with a similarly conserved HisC in Axin1 – as far as we can tell, the only HisC-containing substrate in the human genome capable of co-aggregating with NKD efficiently, implicating Axin1 as its main if not only physiological target. Because of their oligomeric nature, Naked/NKD HisC aggregates attain a high avidity for Axin-HisC, enabling these two proteins to interact efficiently.

The physiological relevance of the HisC-dependent aggregation of Naked/NKD is indicated by specific HisC deletions in human cells: their signaling defects following prolonged Wnt stimulation (reduced β-catenin responses, and stabilization of Axin**; Fig. 5A, B; Fig. S8**) are consistent with precocious termination of Wnt signaling by Axin which, as a result of its stabilization, resumes assembling active β-catenin destruction complexes prematurely. It therefore seems that Naked/NKD acts similarly to the ubiquitin E3 ligase SIAH which targets Axin for proteasomal degradation upon Wnt signaling (Ji et al., 2017). Indeed, the two factors may function redundantly to destabilize Axin, thereby maintaining Wnt signaling responses by preventing Axin from re-establishing quiescence too early.

The marked concentration-dependence of HisC core aggregation (**Fig. 4A**) ensures that Naked/NKD only aggregates once it has accumulated upon prolonged Wnt signaling. Furthermore, aggregation is promoted by Dishevelled, and our evidence indicates that DEP-dependent dimerization of Dishevelled is a prerequisite for facilitating Naked/NKD aggregation. Since the PDZ domain of Dishevelled binds to a dimer of NKD EF-hands, this implies that the interaction between the two proteins results in a considerable increase (>4-fold) in local concentration of Naked/NKD, which may overcome the threshold necessary for triggering HisC aggregation. The dual dependence of Naked/NKD aggregation on Wnt stimulation and Dishevelled dimerization could safeguard against fortuitous spontaneous aggregation and, thus, inappropriate destabilization of Axin.

Given the moderately stable interaction between Dishevelled and Naked/NKD (**Fig. S5**), it seems plausible that the two proteins remain associated following HisC aggregation, which could lead to the destabilization of Dishevelled alongside Axin. In support of this is the stabilization of DVL2 in NKDΔHisC cells during prolonged Wnt stimulation (**Fig. 5B**), which may explain why loss-of-function of Naked/NKD can lead to hyperactive Wnt signaling, and interfere with PCP signaling (Hu et al., 2010; Marsden et al., 2018; Rousset et al., 2001; Schneider et al., 2010; Van Raay et al., 2007; Zeng et al., 2000). Furthermore, it could also explain why overexpressed Naked/NKD downregulates the signaling activity of Dishevelled (Rousset et al., 2001) (**Fig. 1A**) and reduces its levels (Guo et al., 2009; Hu et al., 2010; Schneider et al., 2010). Indeed, it may explain why Naked/NKD loss has little effect on Wnt signaling in some context since its effects in destabilizing Dishevelled (a Wnt agonist) as well as Axin (a Wnt antagonist) simultaneously may cancel themselves out in some circumstances.

Our discovery of the p62 and NBR1 autophagy receptors as Nkd1-interacting proteins suggests that the destabilization of Naked/NKD binding partners is mediated by autophagy. This notion is supported by the pronounced binding preference of p62 to aggregated Nkd1 (**Fig. 8D**), presumably reflecting the high avidity of decameric NKD HisC core aggregates for binding partners. This binding preference implies that the targeting of NKD-binding partners to autophagosomes is contingent on Naked/NKD aggregation. The notion of Naked/NKD targeting its binding partners for autophagy is also consistent with earlier reports that Dishevelled can be degraded by autophagy via binding to p62 (Gao et al., 2010; Ma et al., 2015; Zhang et al., 2011b). Indeed, there appears to be an interaction network between Dishevelled, p62, Axin and aggregated Naked/NKD, implicating Naked/NKD HisC cores as adaptors between Wnt signalosomes and autophagy receptors, as an alternative route to the SIAH pathway (Ji et al., 2017) for the disposal of Axin and other signalosome components upon Wnt signalling. Since this route would involve the envelopment of Naked/NKD co-aggregates by autophagic membranes (Kirkin et al., 2009), it is worth noting that these co-aggregates may not inevitably reach lysosomes because of the endosomolytic properties of highly concentrated histidines (155-265 histidines per single Nkd1-Axin1 co-aggregate): these histidines are expected to become protonated under the mildly acidic conditions typical for endocytic compartments, which could cause swelling and rupture of autophagosomal membranes (Lo and Wang, 2008), thereby potentially allowing escape and recycling of Naked/NKD co-aggregates. This putative endosomolytic activity of Naked/NKD HisC co-aggregates is intriguing, and worth testing in future under physiological conditions.

## MATERIALS AND METHODS

### Plasmid constructions

Mutagenesis of parental plasmid DNA was carried out with standard PCR-based methods, using either KOD DNA polymerase (Merck Millipore), Phusion DNA polymerase (NEB) or Q5 polymerase (NEB), and verified by sequencing. To generate BioID plasmids, the coding sequence for Nkd1 (and mutants thereof) and BirA*(R118G) were amplified by megaprimer PCR and inserted into pcDNA5/FRT/TO using Gibson assembly.

### Cell-based assays

HEK293T cells were cultured in DMEM (GIBCO), supplemented with 10% fetal bovine serum (FBS) at 37°C in a humidified atmosphere with 5% CO_2_. All cells were screened for *Mycoplasma* infection. Transient cell transfections were performed using polyethylenimine or Lipofectamine 2000. Wnt stimulation with Wnt3a-conditioned media was for 6 hrs (unless otherwise stated).

For coIP assays, cells were lysed 24-36 hr post-transfection in lysis buffer (20 mM Tris-HCl pH 7.4, 200 mM NaCl, 10% glycerol, 0.2% Triton X-100, protease inhibitor cocktail and PhosSTOP). DNA amounts were adjusted for equal expression (1 μg HA-Nkd1; 200 ng HA-Nkd1ΔHisC; 300 ng HA-Nkd1ΔEF). Lysates were cleared by centrifugation for 10 minutes (min) at 16,100x *g*, and supernatants were incubated with affinity gel (Flag- or GFP-trap) for 90 min to overnight at 4°C. Subsequently, immunoprecipitates were washed 4x in lysis buffer and eluted by boiling in LDS sample buffer for 10 min.

For SuperTOP assays, cells were lysed <20 hrs post-transfection with SuperTOP and CMV-Renilla (control) plasmids, and analyzed with the Dual-Glo Luciferase Reporter Assay kit (Promega) according to the manufacturer’s protocol. Values were normalized to Renilla luciferase, and are shown as mean ± SEM relative to unstimulated controls (set to 1 in **Figs. 1A & 5A; Figs. S2, S3 & S8**).

### BioID proximity labeling

For BioID experiments, Flp-In HEK293 cell transfections were supplemented with pOG44 recombinase and selected with 250 μg ml^-1^ hygromycin 48 hrs post-transfection. Stable cell lines were induced with 1 μg ml^-1^ tetracycline and treated with Wnt3a-conditioned media and 50 μM biotin 12 hrs prior to lysis. BioID pull-downs were carried out using Streptavidin MyOne Dynabeads essentially as described (Roux et al., 2013), and biotinylated proteins eluted by boiling in LDS sample buffer. Samples were analyzed by SDS-PAGE (BioIP assays) or mass spectrometry (see below).

### Densitometry

Densitometry of Western blots (**Figs. 5A & S8**) was performed on scanned low exposure films using ImageJ software. Where possible, an average of two exposures was taken for each experiment. Values are expressed as arbitrary units obtained for proteins of interest relative to GSK3β as loading controls. For each cell line, values were normalized to its own 0-hour control.

### CRISPR/Cas9 genome editing of human cells

Genome editing by CRISPR/Cas9 of HEK293T cells was essentially as described (van Tienen et al., 2017), using single-guide RNA-encoding plasmid derivatives of pSpCas9(BB)-2A-GFP (PX458; Ran et al., 2013). The CRISPR design tool *crispor.tefor.net* was used to design guide RNAs (**Fig. S7**). Cells were selected for high expression of GFP by fluorescence-assisted cell sorting 48 hrs post-transfection, and individual clones expanded in 96-well plates. Individual cell clones were screened by sequencing (**Table S3**) to confirm the presence of frameshifting lesions. To ensure consistency and to guard against off-target effects, multiple lines were isolated and characterized in each case.

### CRISPR/Cas9 genome editing in Drosophila

Single-guide RNA sequences (designed using *crispor.tefor.net;* **Fig. S9**) were inserted into pCFD3 by *Bbs*I digestion, as previously described (van Tienen et al., 2017). Embryos from *nos.cas9* flies (from Simon Bullock) were collected for 1 hr and dechorionated in 8% sodium hypochlorite solution for 2 min before injection with DNA (200 ng µl^−1^) under 10S oil. For genotyping, DNA was extracted from individual flies by standard methods, and sequences were determined after PCR amplification (**Table S3**).

### Analysis of embryonic cuticles

Embryos were collected overnight and staged for 24 hrs at 25°C. Unhatched embryos were dechorionated in 8% sodium hypochlorite for 3 min, washed in water and vortexed for 1 min in 1:1 heptane:methanol (−20°C) to remove the vitelline membrane. Embryos from the bottom of the tube were transferred onto a slide and mounted in 1:1 lactic acid:Hoyer’s-based medium (Hoyer’s mountant), and slides were incubated at 65°C overnight. Imaging of cuticles was done with a Zeiss Axiophot microscope (camera Zeiss AxioCam MRc5). Images were taken with differential interference contrast (DIC) optics with a 100x objective (**Figs. 7B-D; Figs. S10-S12**), or dark-field illumination with a 10x objective (**Figs. 6A-F & S11**). For the quantification of denticle numbers (**Fig. 7B, C**), near-normal-looking abdominal denticle belts were chosen under dark field, and denticle numbers were counted at high magnification; for **Fig. 7B**, belts from abdominal segments 3-6 were chosen.

### Antibody staining of embryos

Dechorionated embryos were washed in water and fixed by shaking in 1:3 phosphate-buffered saline (PBS) and heptane containing 4% formaldehyde for 30 min. Ice-cold methanol was added, and tubes were vortexed for 1 min. Embryos were washed in methanol 5x and rehydrated by 5 min washes in 90%, 75%, 50%, 25% of methanol followed by a final PBS wash. Embryos were permeabilized in PBT (PBS containing 0.1% Triton X-100) for 30 min, blocked in BBT (1% BSA in PBT) for 1 hr and incubated with primary antibodies (α-β-gal 1:500; α-Wg 1:10; α-En 1:10) overnight at 4°C. After washing (3x in BBT), embryos were incubated with biotinylated goat-anti-mouse IgG antibody (1:100 in BBT) for 4 hrs and washed (2x in BBT, 1x in PBS). Detection was carried out with ABC Elite kit according to the manufacturer’s protocol. Homozygous *nkd*^*13*^ embryos were identified by the lack of *twi.lacZ*-bearing balancer chromosomes.

### Mass spectrometry

Mass spectrometry was performed by the LMB Mass Spectrometry Facility. Briefly, peptides from *in situ* trypsin digestion were extracted in 2% formic acid/2% acetonitrile mix. Digests were analyzed by nano-scale capillary LC-MS/MS using an Ultimate U3000 HPLC and C18 Acclaim PepMap100 nanoViper (Thermo Scientific Dionex). LC-MS/MS data were searched against a protein database (UniProt KB) with the Mascot search engine program (Matrix Science). MS/MS data were validated using the Scaffold program (Proteome Software). MALDI-TOF mass spectrometric measurements were carried out in positive ion mode on an Ultraflex III TOF-TOF instrument (Bruker Daltonik), using sinapinic acid (Sigma) as matrix.

### Protein expression and purification

Protein-coding sequences were inserted in pET vectors bearing His6-Lip (PDZ: residues 285-373 from human DVL2), His6-RFP (EF-hand: residues 110-190 from human NKD1), Strep-MBP and Strep-Lip (HisC sequences; **Fig. 3A**). Proteins were expressed in *E. coli* BL21DE3 pRare2 cells overnight at 24°C after induction with IPTG at OD600 of 0.7-0.9. Cells were pelleted for 20 min at 7000 g, resuspended in 30 mM Tris-HCl, 300 mM NaCl, pH 7.4 and lysed with sonification (10 sec on, 10 sec off, 2 min in total, 90% amplitude). Lysates were cleared by centrifugation at 100 000x *g* for 20 min, and clear supernatant was loaded onto NiNTA beads. Beads were washed with 10 column volumes of 30 mM Tris-HCl, 300 mM NaCl, pH 7.4, and proteins were eluted in the same buffer, supplemented with 300 mM imidazole. Fractions containing protein of interest were pooled and further purified by SEC with a S200 Superdex 26/60 column in 15 mM Tris-HCl, 150 mM NaCl, pH 7.4, or in 15 mM acetate, 150 mM NaCl, pH 5.0.

### ITC

ITC was performed on MicroCal iTC200 in 15 mM Tris-HCl, 150 mM NaCl, pH 7.4 by injecting Lip-DVL2-PDZ or Lip-DVL2-PDZ-V334E (696 μM) into 17.2 μM solution of RFP-NKD-EF (preinjection of 0.5 μL + 19 injections of 2 μL). The data were analyzed following the manufacturer’s guidelines.

### SEC-MALS

Purified proteins and preformed complexes were analyzed with an Agilent 1200 Series chromatography system connected to a Dawn Heleos II 18-angle light-scattering detector combined with an Optilab rEX differential refractometer (Wyatt). Samples were loaded onto a Superdex-200 10/300 gel filtration column (GE Healthcare) at 2 mg ml^-1^ and run at 0.5 mL/min in buffer (15 mM TrisHCl, 150 mM NaCl, pH 7.4), and the data were processed with Astra V software.

### Co-aggregation assay

Protein samples were prepared in 15 mM acetate, 150 mM NaCl, pH 5.0 and mixed in different ratios. After 10 min, 1 M Tris-HCl, pH 8.0 was added to a final concentration of 200 mM. After 20 min incubation, LDS sample buffer was added, and samples were boiled for 10 min.

### Quantitation and statistical analysis

All error bars are represented as mean ± SEM from >3 independent experiments. Statistical significance was calculated by one-way ANOVA with multiple comparisons (**Figs. 1A & S3**), two-way ANOVA with multiple comparisons (**Figs. 5A, B & S8**) or Student’s t tests (**Figs. 7B & S2**), and denoted as follows: ^*^ = p < 0.05, ^**^ = p < 0.01, ^***^ = p < 0.001, ^****^ = p < 0.0001.

## AUTHOR CONTRIBUTIONS

M. G. and J. E. F. performed the cell-based assays, M. R. carried out the biochemical analysis with recombinant proteins, J. M. conducted the genetic analysis in *Drosophila*, and M. B. conceived and supervised the study, and wrote the manuscript with input from all authors.

## ACKNOWLEDGEMENTS

We thank Keith Wharton for plasmids, Mark Skehel and his team for mass spectrometry, Maria Daly for cell sorting, Balaji Santhanam for advice on the bioinformatics, Chris Johnson for technical assistance with the biophysics, and Hugh Pelham for discussion. This work was supported by Cancer Research UK (C7379/A24639) and the Medical Research Council (U105192713).

## FIGURE LEGENDS

**Fig. S1 (related to main Fig. 1).**
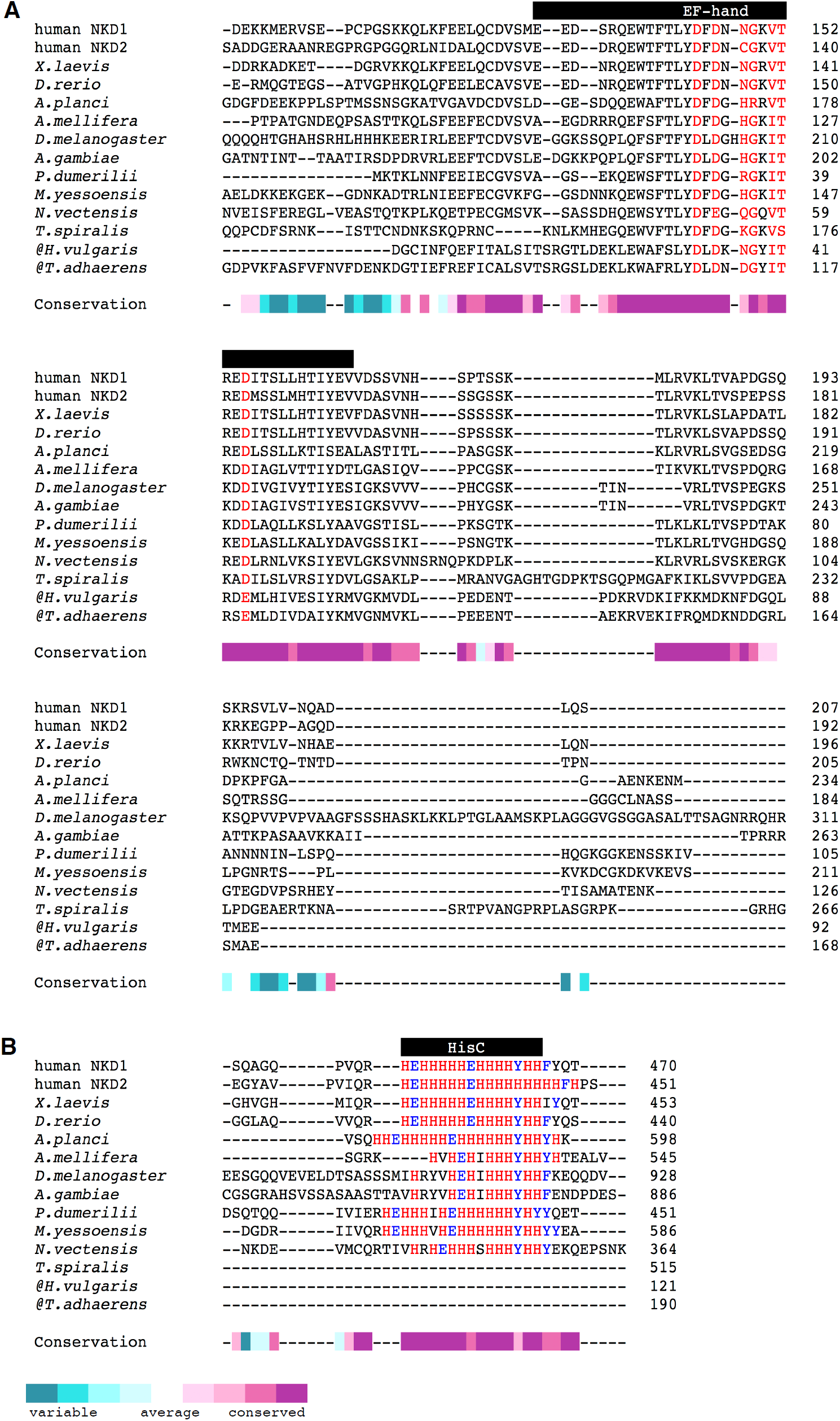
Sequence conservation of Naked/NKD orthologs. Sequence alignments of the (**A**) EF-hand and the (**B**) C-terminal HisC cluster across diverse animal species. @, C-terminal sequences not available. Conservation score was calculated with the ConSurf webserver.

**Fig. S2 (related to main Fig. 1).**
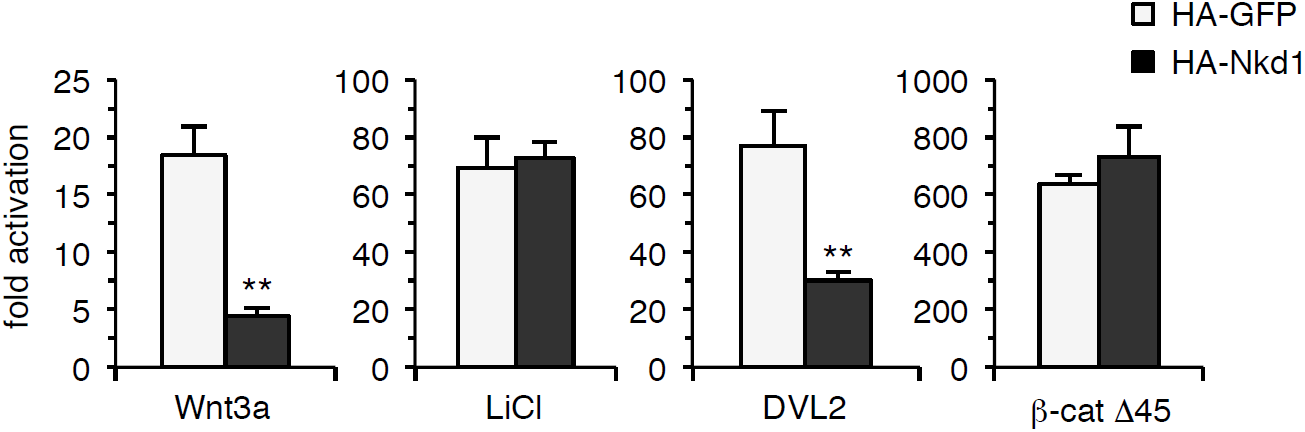
Effects of overexpressed Nkd1 on β-catenin signaling. SuperTOP assays, monitoring the effects of overexpressed HA-Nkd1 on β-catenin-dependent transcription in response to stimulation by Wnt3a or LiCl (6 hours), or to overexpressed DVL2-GFP or activated β-catenin (β-cat Δ45) in transfected HEK293T cells; error bars, SEM of >3 independent experiments; Student’s t-test; ** = p < 0.01.

**Fig. S3 (related to main Fig. 2).**
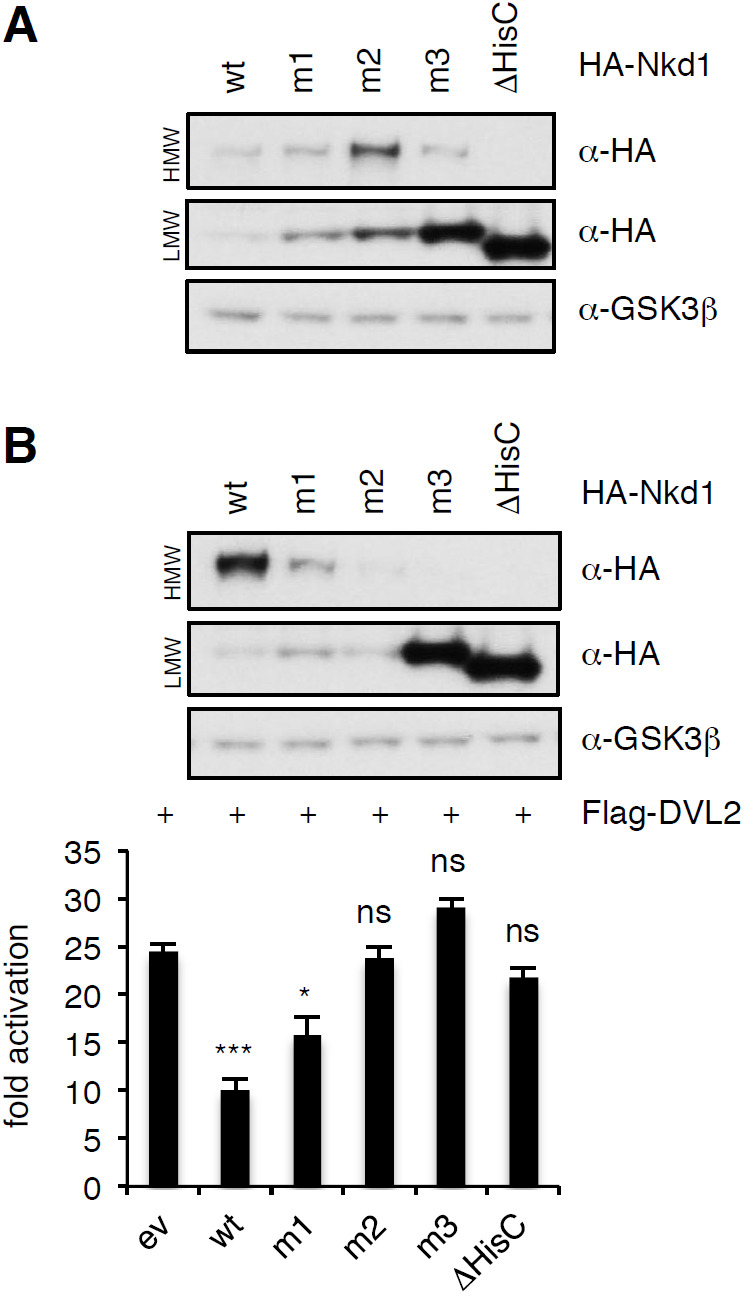
Effects of overexpressed Nkd1 HisC mutants on β-catenin signaling. Western blots monitoring expression of HA-Nkd1 and specific HisC point-mutants thereof (see **Fig. 2D**) alone (**A**) or in the presence of Flag-DVL2 (**B**). SuperTOP assays in transfected HEK293T cells, monitoring the effects of overexpressed HA-Nkd1 on β-catenin-dependent transcription owing to overexpressed Flag-DVL2; error bars, SEM of >3 independent experiments; one-way ANOVA with multiple comparisons; * = p < 0.05, ^***^ = p < 0.001, ns = not significant.

**Fig. S4 (related to main Fig. 3).**
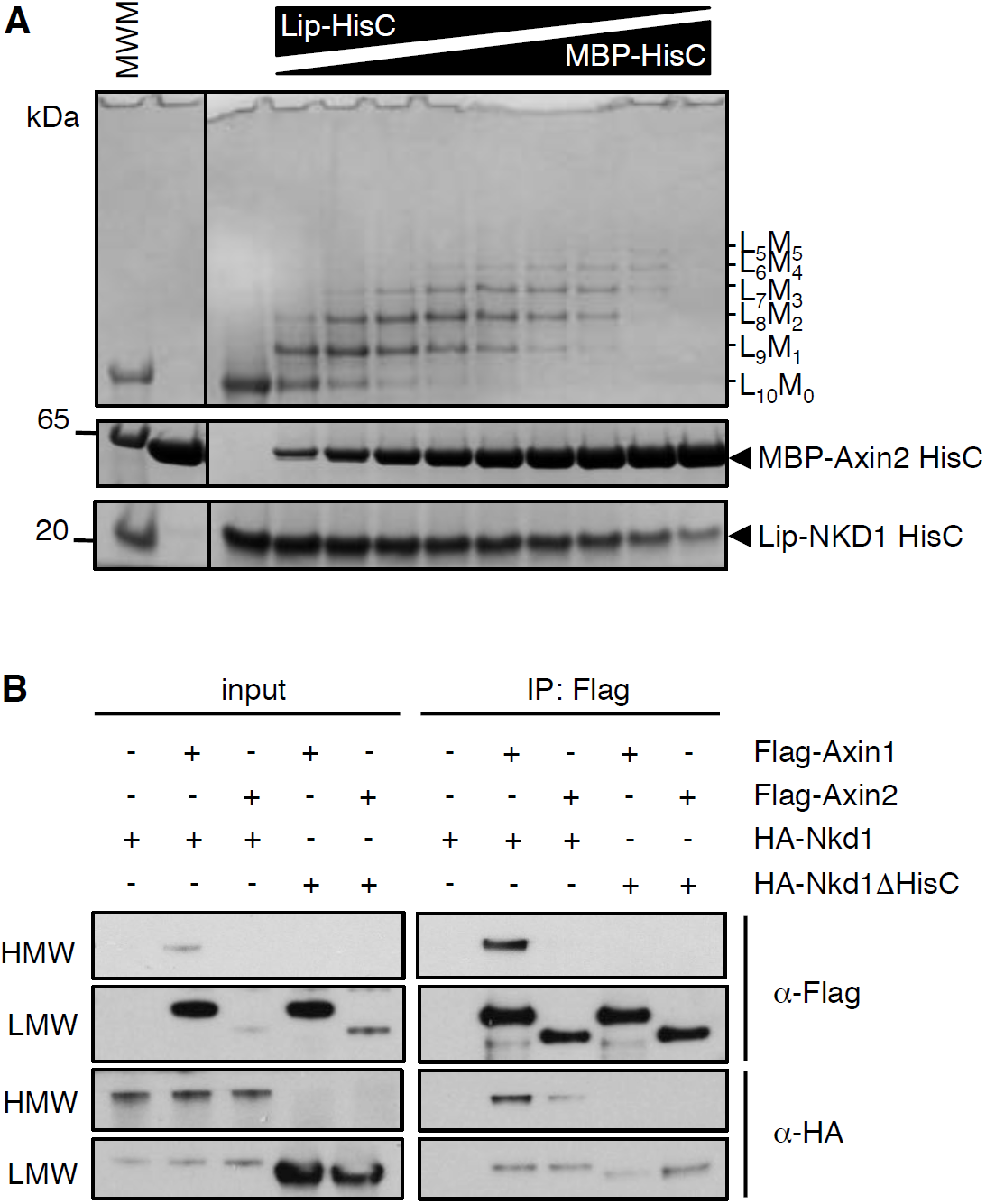
Co-aggregation between NKD1 and Axin2. (**A**) Co-aggregation assays after mixing purified recombinant Lip-NKD1-HisC and MBP-Axin2-HisC (KHVHHHYIHHHA) at different ratios (MWM, molecular weight markers). (**B**) CoIP assays between proteins co-expressed in transfected HEK293T cells, as indicated above panels, revealing formation of HMW co-aggregates between HA-Nkd1 and Flag-Axin1 but not Flag-Axin2.

**Fig. S5 (related to main Fig. 3).**
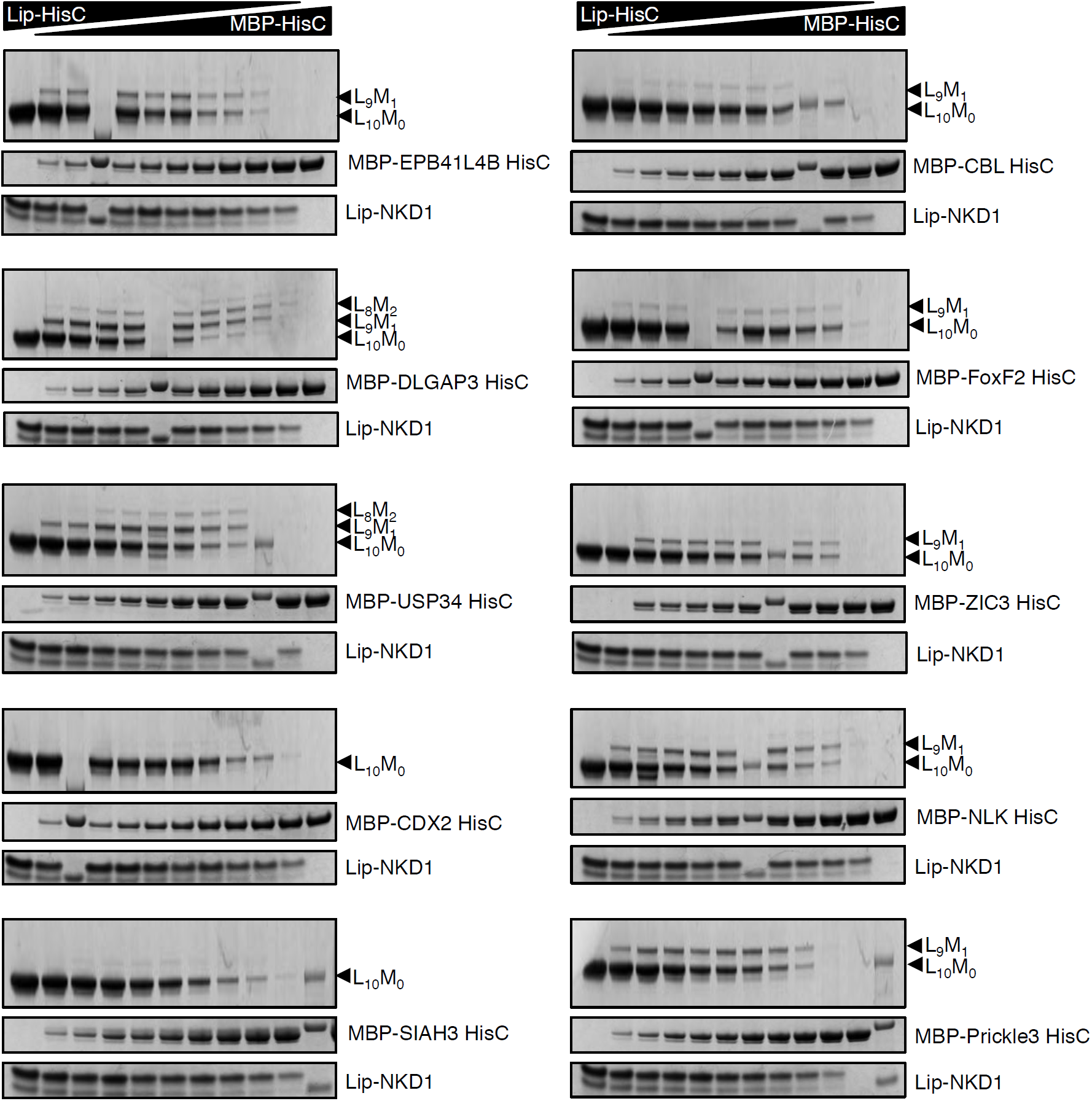
Systematic co-aggregation tests with NKD1-HisC. Co-aggregation assays after mixing purified recombinant Lip-NKD1-HisC and MBP-HisC from selected proteins, as indicated on the right (see main **Fig. 3A**, for HisC sequences).

**Fig. S6 (related to main Fig. 4).**
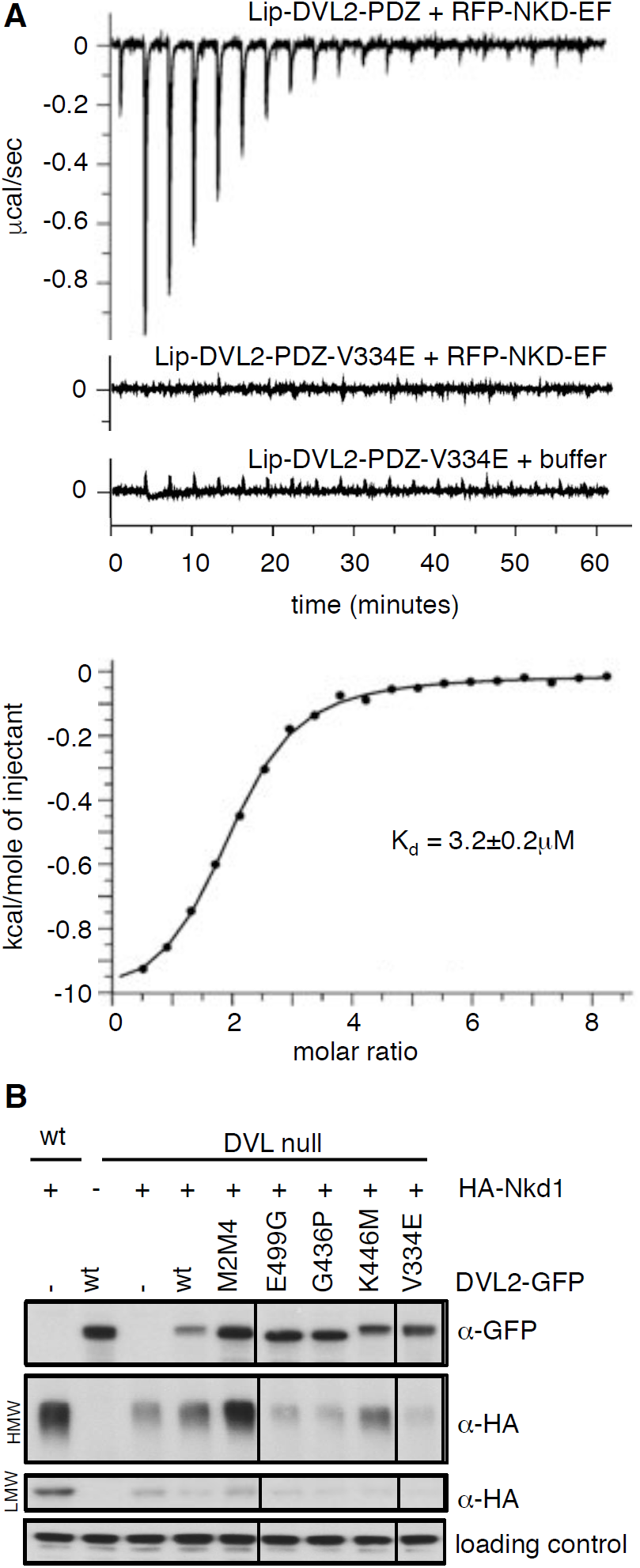
Binding of the NKD1 EF-hand to the DVL2 PDZ cleft. (**A**) ITC profiles for RFP-NKD1-EF binding to Lip-DVL2-PDZ or V334E cleft mutant; *K*_d_ value, Lip-DVL2-PDZ binding to RFP-NKD1-EF. (**B**) Western blots monitoring the formation of Nkd1 HisC aggregates dependent on complementation of DVL null cells by re-expression of wt or mutant DVL2-GFP (see also main **Fig. 4D**).

**Fig. S7 (related to main Fig. 5).**
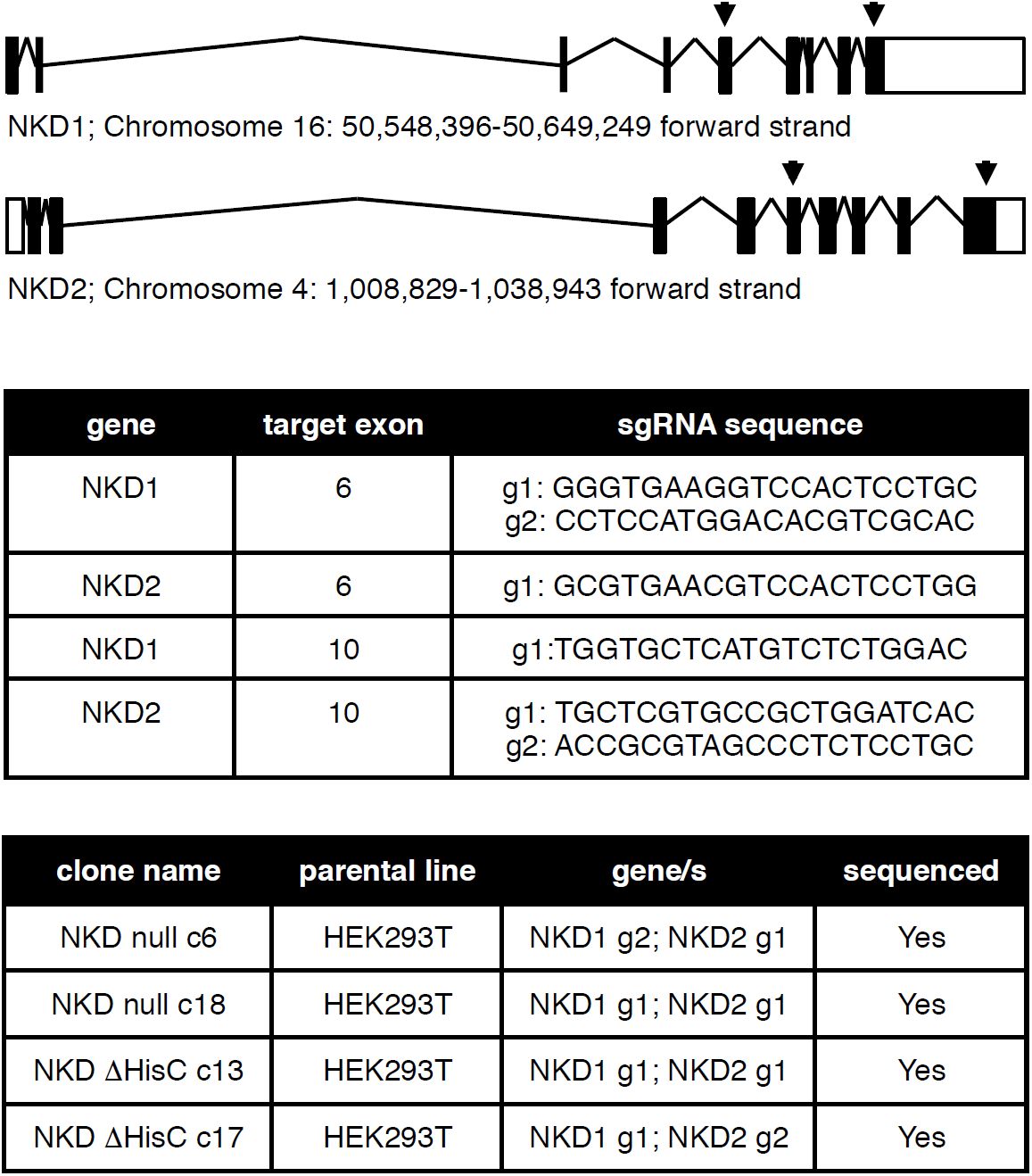
CRISPR engineering of HEK293T cells. *Top*, genomic locations and organizations of the human *NKD1* and *NKD2* loci, with positions of CRISPR gRNAs indicated; *underneath*, sequences of gRNAs used for generating truncations in two different NKD null (c6 and c18) or ΔHisC (c13 and c17) lines, each verified by sequencing of both genes.

**Fig. S8 (related to main Fig. 5).**
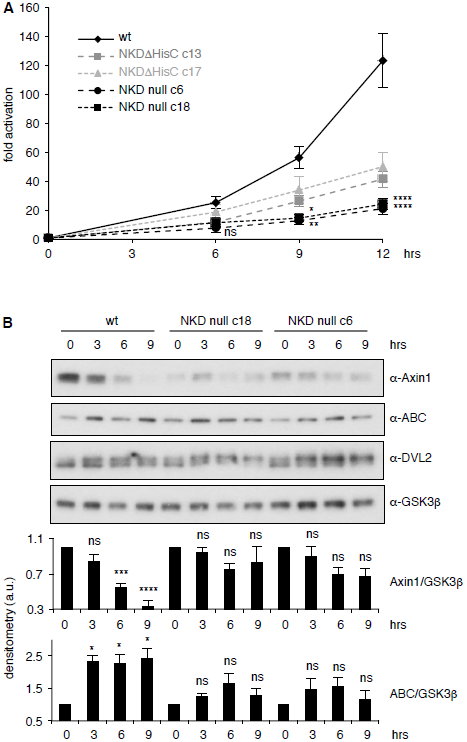
NKD is required for sustained Wnt signaling. (**A**) SuperTOP assays as in main **Fig. 5A** (same values), with statistical tests (two-way ANOVA with multiple comparisons) for c6 and c18 null lines; * = p<0.05, ** = p<0.01, **** = p<0.0001 (**B**) Western blots corresponding to SuperTOP assays shown in main **Fig. 5A**, monitoring expression levels of endogenous proteins in NKD null cells, quantified by densitometry underneath, normalized to 0 hr time point for each cell line (a.u., arbitrary units); SEM of >3 independent experiments; two-way ANOVA with multiple comparisons; * = p<0.05, *** = p<0.001, **** = p<0.0001; ns = not significant. We note that the levels of endogenous Axin1 are consistently reduced in unstimulated NKD null cells compared to their parental controls, perhaps reflecting a compensatory mechanism of these null cells to maximize their tonic Wnt signaling levels (see also Gammons et al., 2016b).

**Fig. S9 (related to main Fig. 6).**
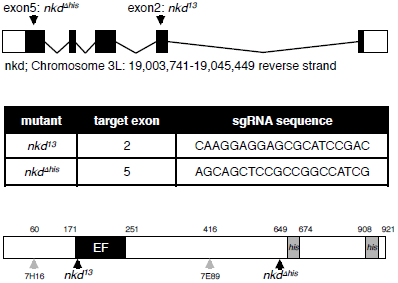
CRISPR-engineered Drosophila *nkd* alleles. *Top*, genomic location and organization of the *Drosophila nkd* locus, with positions of CRISPR gRNAs indicated; *underneath*, sequences of gRNAs used for generating two different *nkd* truncations, each verified by sequencing; *bottom*, cartoon of Naked, with domains and positions of various alleles indicated.

**Fig. S10 (related to main Fig. 6).**
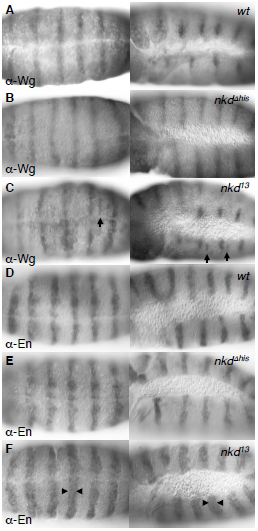
Ectopic Engrailed and Wingless expression in *nkd* mutant *Drosophila* embryos. 5-6 hrs old wt and *nkd* mutant embryos, as indicated in panels, stained for Wingless (Wg, **A-C**) or Engrailed (En, **D-F**) expression; *left*, ventral view; *right*, side view. *nkd* null (*nkd*^*13*^) but not ΔHisC mutants (*nkd*^*Δhis*^) exhibit an expansion of En expression (arrowheads) in every segment, and sporadic patches of ectopic Wg expression (arrows). The latter appear to correlate with the sporadic regions of excess naked cuticle shown in main **Fig. 6D, E**, and thus likely accounts for the partially ‘naked’ phenotype of these embryos.

**Fig. S11 (related to main Fig. 6).**
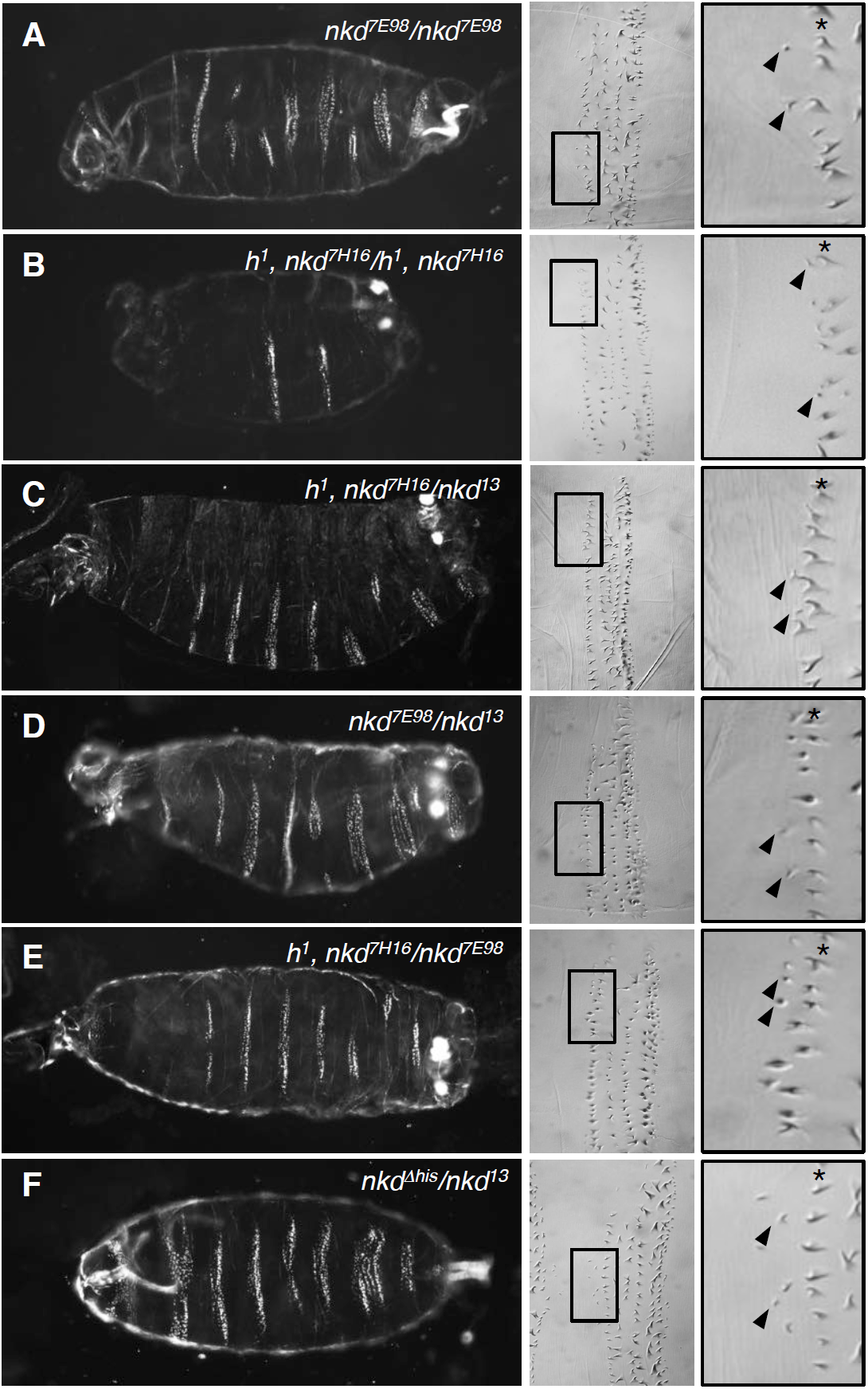
Excess anterior denticles in transheterozygous embryos bearing different combinations of *nkd* alleles. Images of ventral cuticles of embryos bearing different combinations of *nkd* and *h*^*1*^ alleles as indicated in panels, exhibiting excess anterior denticles. Note that the dark field images (on the left) are representative examples for each genotype, while the DIC images (on the right) show selected near-normal denticle belts with excess row-0 denticles (arrowheads); asterisks (*) indicate row-1 denticles.

**Fig. S12 (related to main Fig. 6).**
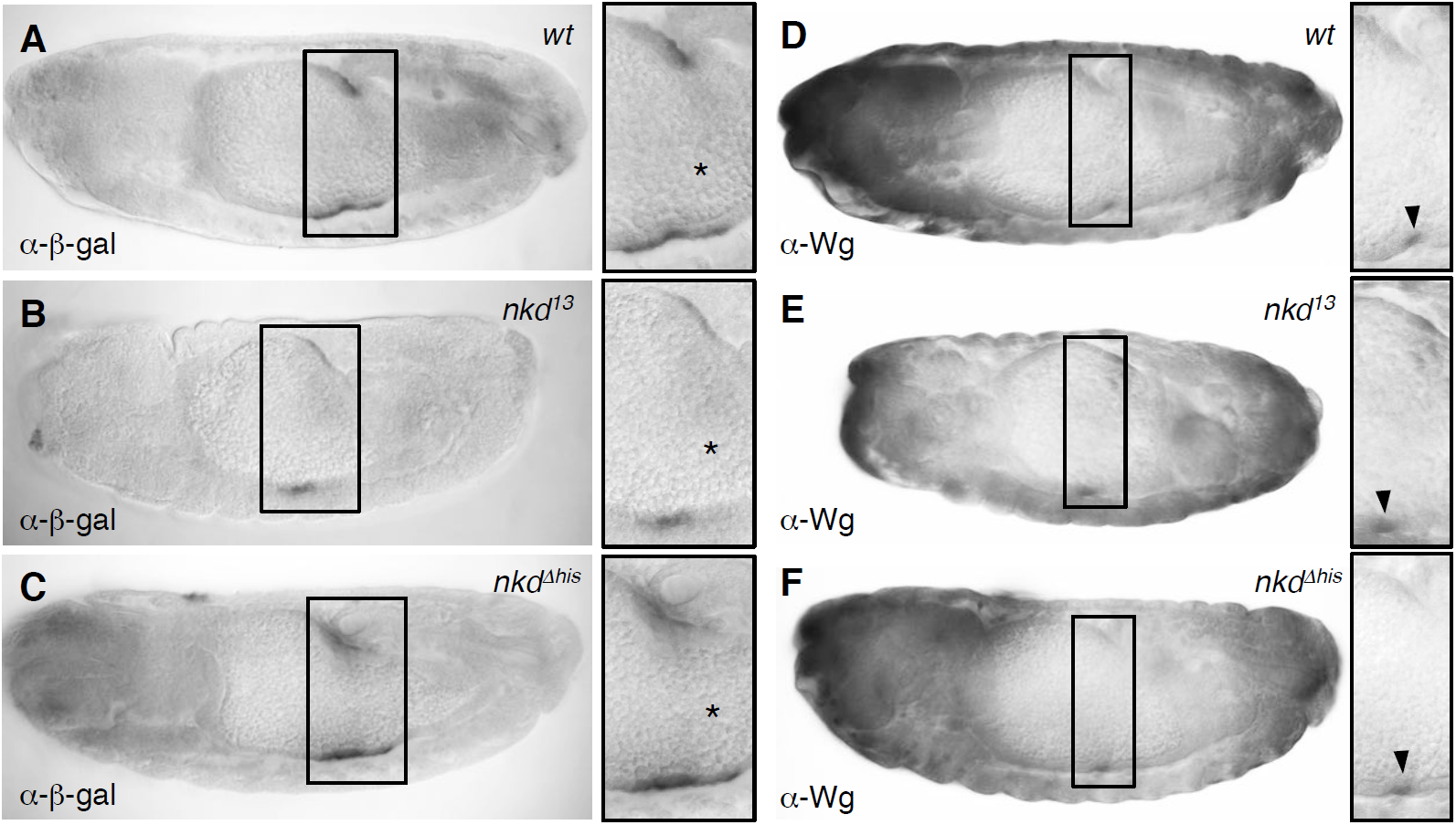
Reduced expression of Wg-responsive *Ubx* midgut reporter gene in *nkd* null mutant embryos. (**A-C**) Side views of 12-14 hrs old embryos bearing *UbxB.lacZ*, fixed and stained with α-β-galactosidase (α-β-gal) antibody, showing reduced expression of *UbxB.lacZ* in *nkd*^*13*^ but not *nkd*^*Δhis*^ homozygous embryos (* in high-magnification views mark positions of Wg signaling sources in the middle midgut). (**D-F**) Side views of −12 hrs old wt or *nkd* mutant embryos, fixed and stained with α-Wg antibody, revealing apparently normal Wg expression (marked by arrowheads in high-magnification views) in the middle midgut of *nkd* mutants.

**Table S1 (related to main Fig. 3).**
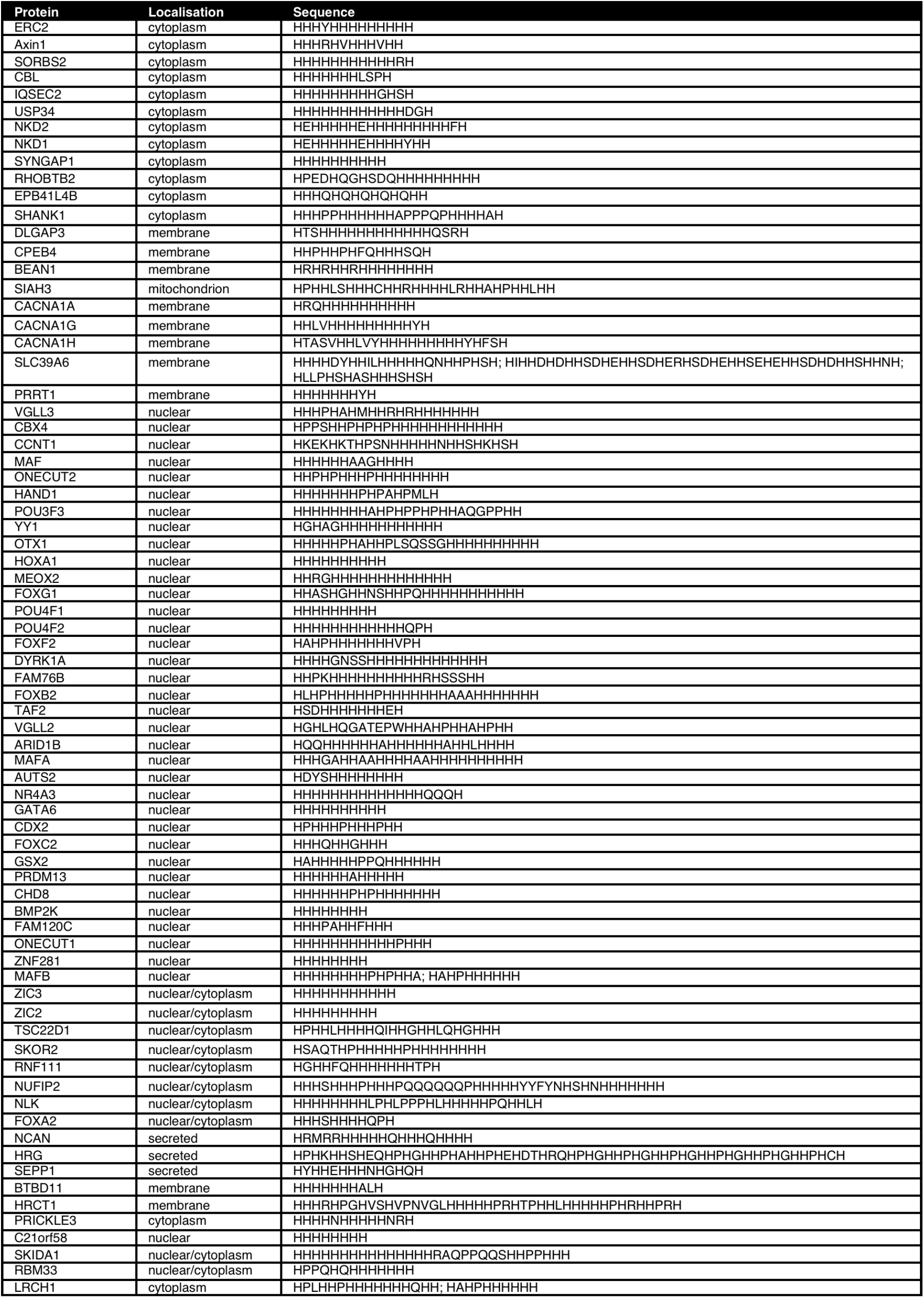
HisC proteins in the human genome and their subcellular location.

**Table S2 (related to main Fig. 5).**
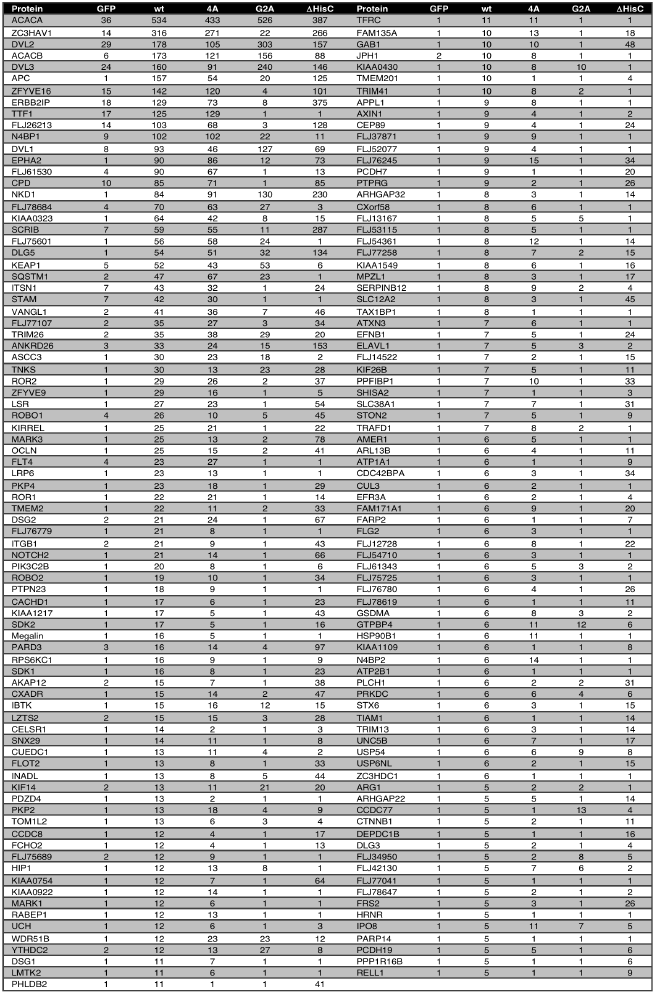
BioID hits obtained with wt or mutant Nkd1-BirA*.

**Table S3 (related to STAR Methods).**
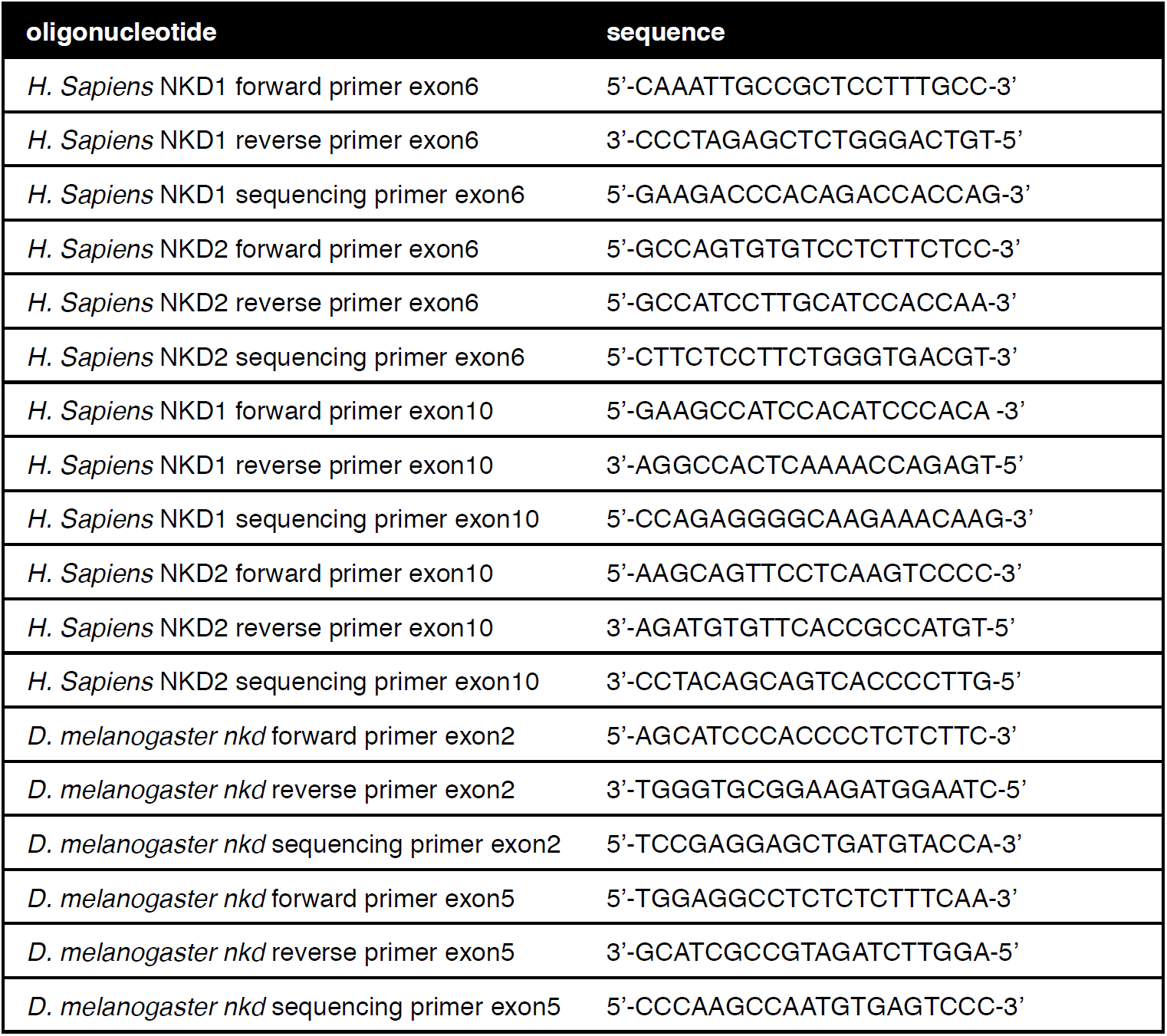
Oligonucleotides used for CRISPR engineering of *NKD1* and *NKD2* loci in HEK293T cells, and of *Drosophila nkd*, for confirmation of CRISPR-engineered lesions.

